# Biocomposite Thermoplastic Polyurethanes Containing Evolved Bacterial Spores as Living Fillers to Facilitate Polymer Disintegration

**DOI:** 10.1101/2023.10.10.561602

**Authors:** Han Sol Kim, Myung Hyun Noh, Evan M. White, Michael V. Kandefer, Austin F. Wright, Debika Datta, Hyun Gyu Lim, Ethan Smiggs, Jason J. Locklin, Md Arifur Rahman, Adam M. Feist, Jonathan K. Pokorski

## Abstract

The field of engineered living materials (ELMs) seeks to pair living organisms with synthetic materials to generate biocomposite materials with augmented function since living systems can provide highly-programmable and complex behavior. ELMs have typically been fabricated using techniques in benign aqueous environments, limiting their application. In this work, biocomposite fabrication was demonstrated in which spores from polymer-degrading bacteria were incorporated into a thermoplastic polyurethane (TPU) using high-temperature melt processing. Bacteria were engineered using adaptive laboratory evolution to improve their heat tolerance to ensure nearly complete cell survivability during manufacturing at 135 °C. Furthermore, the overall tensile properties of spore-filled TPUs were substantially improved, resulting in a significant improvement in toughness. The biocomposites facilitated disintegration in compost in the absence of a microbe-rich environment. Finally, spores retained a programmed function, expressing green fluorescent protein. This research provides a scalable method to fabricate advanced biocomposite materials in industrially-compatible processes.

## 1. Introduction

Engineered living materials (ELMs) is a burgeoning field in which living and synthetic matter are combined to provide composite materials with augmented and complex functions, far beyond what a traditional polymeric material could accomplish alone. The promise of incorporating living matter into biocomposites has generated materials that are capable of responding to stimuli^1^ (i.e. light, nutrients, inducers, etc), and consequently morphing their shapes^2,3^ and/or properties^4^, which have been utilized for living biosensors^5^, wearable bioelectronics^5,6^, drug delivery systems^7^, wound healing patches^8^ and self-regenerating skin^9^. Introducing live cells into polymer composites as a sustainable and ‘smart’ filler material has the potential to greatly improve both material properties and their ecological footprint. Live cells have ideal features as smart polymer additives such as self-replication, self-regulation, and programmable stimuli-responsiveness^10^. Moreover, cells can be genetically programmed to synthesize both small and large molecules and can be further engineered to render other diverse functionalities^11^. Successfully harnessing living cells has limitless potential to develop polymer composites with enhanced properties such as improved mechanical performance and other performance characteristics, such as programmed/accelerated disintegration. Despite this potential, live cells have rarely been exploited as polymer additives in practice due to their fragility. Cells require careful handling in terms of hydration, osmotic pressure, temperature and pH, compared to other non-living biological substances. Polymer processing into commercial parts typically requires heat, shear stress and/or solvents, all of which are detrimental to cell viability. To that end, incorporating live cells into polymers has only been demonstrated with limited types of polymers, which can be fabricated under mild conditions (i.e. low melting temperature, aqueous conditions, and low shear stress)^1,8,11,12^, and in most studies it is unclear if the cells maintained viability. To render cell-based biocomposite materials broadly useful, the poor stability of living cells must be resolved to use them in industrially-relevant fabrication processes.

Some bacteria have naturally evolved to resist extreme conditions by forming spores^13^. Spores can preserve their viability against high temperatures, pressure, toxic chemicals (i.e. acids, bases, oxidants, and organic solvents), and radiation^14,15^. The robustness of spores is attributed to multiple factors that protect chromosomal DNA in the core, such as multi-layered protein coats, a peptidoglycan cortex and a partially dehydrated core crowded with protective intracellular molecules^16,17^. Bacterial spores are metabolically dormant yet can survive for years. Even though spores have minimal metabolic activity, they are poised for germination into vegetative cells within minutes^18–20^.

*Bacillus* s*ubtilis* are one of the most well-known spore-forming bacteria and have unique metabolic properties. They are ubiquitous in nature and have potential health benefits as FDA-approved probiotics^21–23^. Some *B. subtilis* strains have degradation activity toward polyester-based polymers^24,25^. Collectively, the high stability, safety, polyester degradation activity, and triggerable sporulation/germination of *B. subtilis* make them promising additives in developing biocomposite polymers with programmed biodegradation. However, a significant challenge to employ *B. subtilis* spores in industrially-relevant polymer processing is their lack of heat tolerance. Spores of several *B. subtilis* strains are known to be resistant at ∼ 100 °C for several minutes^26^, but most industrial thermoplastic processing requires higher temperatures above 130 °C^27^. In fact, *B. subtilis* lost >90% of spore viability within 1 minute at these temperatures^28,29^.

Adaptive Laboratory Evolution (ALE) is an evolutionary engineering approach that has been effectively adopted in situations where the genetic causality to increase a phenotypic property is not defined or intuitive^30–32^. By capitalizing on the naturally occurring mutation during growth and division, coupled with growth-based selection, ALE effectively enhances desired phenotypes. This versatile approach finds wide applicability across various microorganisms that can be cultured in a laboratory setting. Notably, ALE has been effective for optimizing *Bacillus* species specifically in the past for enhancing tolerance towards inhibitors in hydrolysed biomass^33^, as well as stress tolerance to low pressures^34,35^, and establishing a non-native pathway^36^. Thus, ALE is a well-suited strategy to augment the heat tolerance of *Bacillus* spores^37,38^, thereby enabling their use as a functional living additive material.

We envisioned that the combination of evolutionarily-engineered *B. subtilis* spores and polyester-based thermoplastic polyurethanes (TPUs) would have synergistic effects in both improving the mechanical properties and programming/facilitating the degradation of a spore-filled biocomposite TPU. The multilayered proteins found in the exospore contain complex macromolecular structures which contribute to the glass transition temperature and, thus, the elastic moduli of bacterial spores^39^. Therefore, spores can simultaneously serve as particulate soft or rigid fillers to reinforce a TPU matrix, as well as biocatalysts for TPU disintegration under their dormant and germinated forms, respectively (**Fig. 1**). We demonstrated that heat-shock tolerized *B. subtilis* spores retained ∼100% viability in TPUs after hot melt extrusion (HME). HME is the most widely-used and industrially-scalable technique in the polymer industry and forms the basis for most polymer manufacturing technology^40,41^. The resulting biocomposite TPU showed significantly improved tensile properties, when compared to neat TPU without spores or to non-evolved spore composites. Spores in the TPU matrix could be germinated by nutrients in compost and germinated *B. subtilis* cells improved the kinetics of end-of-life TPU disintegration in compost regardless of the microbial activity/diversity. Lastly, biological function was successfully programmed into the biocomposite material by genetically engineering *B. subtilis* to express a model green fluorescent protein (GFP). Plasmids transformed into *B. subtilis* were retained in spores after HME, and fluorescence could be easily detected once the spores were germinated. Overall, this work presents a scalable method for the fabrication of biocomposite materials with improved mechanical properties and programmed biological functionalities.

**Fig. 1.**
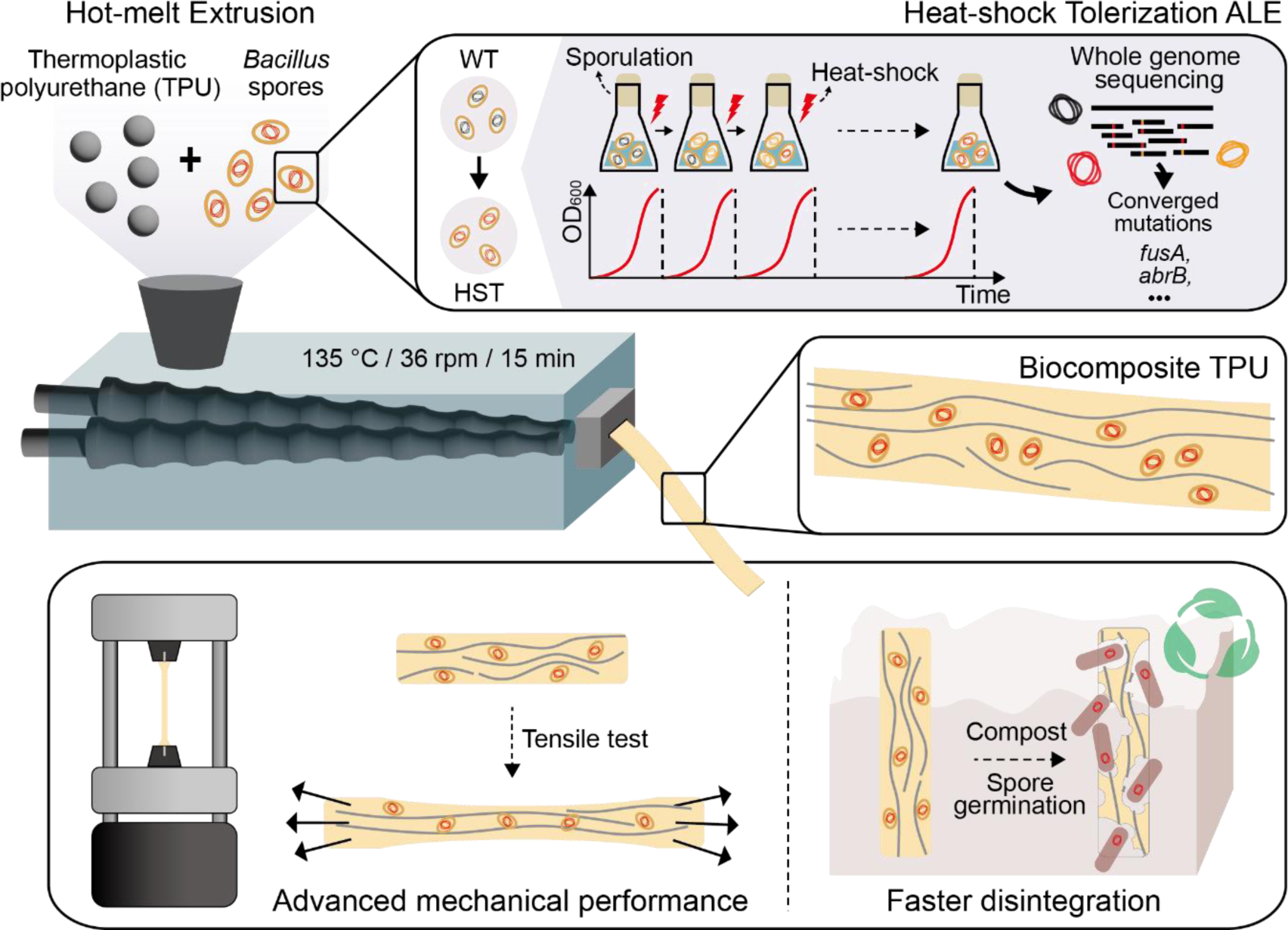
Schematic representation of the fabrication of a biocomposite TPU filled with bacterial spores (WT; wild-type ATCC 6633, HST; Heat-shock tolerized strain) capable of degrading TPUs. Heat-shock tolerance of spores was enhanced through the Adaptive Laboratory Evolution (ALE). The biocomposite TPU was fabricated via hot melt extrusion, wherein dormant spores acted as particulate fillers reinforcing the mechanical properties of TPU matrix. During the end-of-life cycle testing, germinated bacterial cells enhanced disintegration of the TPU.

## 2. Results and Discussion

### 2.1. Adaptive Laboratory Evolution of *Bacillus* spores to increase heat-shock tolerance

To identify *Bacillus* strains with inherent TPU degradation and assimilation capabilities^42^, we evaluated the growth of five common *Bacillus* strains (American Type Culture Collection; ATCC 23857, ATCC 6051, ATCC 7061, ATCC 21332, ATCC 6633) in minimal media supplemented with TPU powder (ground Elastollan^®^ BCF 45 gifted from BASF) as the sole carbon and energy source. Among the candidates, the ATCC 6633 strain exhibited robust growth and efficient assimilation of TPU powder, making it a suitable host strain for further study (**Supplementary Fig. 1**).

Previous research has shown that *Bacillus* spores^26^ display robust heat-shock resistance but also exhibit differences in thermal tolerance across species. The ATCC 6633 strain has demonstrated significant heat tolerance in dry-heat conditions^43^, however, we deemed this to be insufficient for melt processing and pursued an ALE experiment to further promote thermal stability of spores. We first characterized the initial heat-shock tolerance of spores by exposing them to boiling water (100 °C, i.e., wet-heat) for varying lengths of time (**Fig. 2A**) and the germination efficiency (colony forming units, CFU) of the treated spores compared to the untreated spores was assessed. Although the ATCC 6633 spores exhibited considerable heat tolerance in dry-heat conditions over 120 °C^26^, exposure to wet-heat of 100 °C for even a few minutes resulted in a sharp decrease in viability. Only 24.7% of spores were able to germinate after 1 minute of heat treatment, which further decreased to 3.3% after 3 minutes. TPU melt processing is performed at an even higher temperature indicating the heat-shock tolerance is imperative for biocomposite fabrication.

**Fig. 2.**
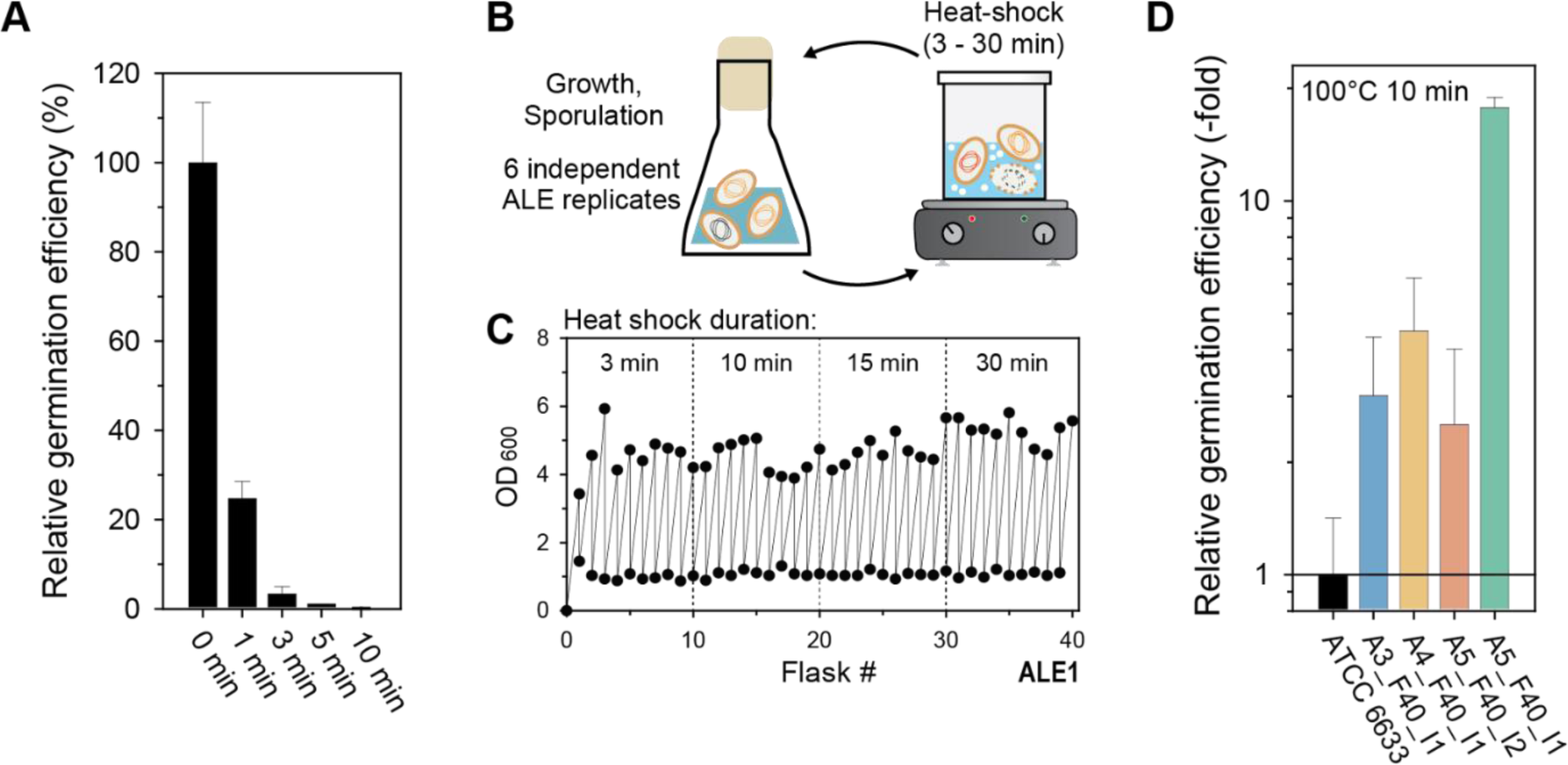
(A) Characterization of initial heat-shock tolerance of Bacillus subtilis ATCC 6633 spores. Germination efficiency was measured after exposure to boiling water for different durations (minutes). (B) Conceptual design of an ALE experiment for improving heat-shock resistance of spores. (C) Representative growth profiles of one replicate of the ALE experiment (D) Characterization of enhanced heat-shock tolerance for ALE-derived clones after exposure to boiling water for 10 mins. Error bars indicate standard deviations from three independent experiments.

To enhance the heat-shock tolerance of *B. subtilis* spores, ALE experiments were conducted (**Fig. 2B-C**). Spores were generated after 24 h of cell culture in Difco sporulation medium (DSM) and spore cultures were subjected to boiling water and then propagated into fresh DSM media (**Fig. 2B**). The initial heating time was set to 3 min, which was enough to render >90% of spores dead. Subsequent heat tolerance passages were gradually increased to 30 min during the ALE experiments (**Fig. 2C**). Cell densities (OD_600_) were measured at the start and after 24 hours of culture to confirm sporulation. The ALE experiments were conducted for 40 cycles and were parallelly implemented with 6 independent lineages all from the same starting genotype to examine adaptive convergence^44^. All six ALE end-point lineages could continuously produce spores after 30 min of heat treatment (**Fig. 2C**), whereas the wild-type starting strain spore cultures could not show any detectable increase in cell density. This finding indicated the ALE approach was effective in selecting for strains with enhanced heat-shock tolerance properties when in spore form.

### 2.2. Identification of the genetic basis for enhanced heat-shock tolerance of *Bacillus* spores

To identify the causal mutations enabling the increased heat-shock tolerance of the ALE-derived strains, whole-genome sequencing was conducted for populations and included two isolates from each of the final flasks of each ALE lineage (n = 6) (**Supplementary Data 1**). Two frequently mutated genes, *fusA* and *abrB*, were most prevalent across the parallel ALE lineages and were the focus of mutation causality assessment (**Table 1**). Mutations affecting these two genes made up 8 out of a total of 18 unique mutations (44.4%) identified from all samples (**Supplementary Data 1**). Overall, there was an average of 3 mutations per sample and a mode of 2 mutations for both population and clonal samples (lineages 4 and 6 had identical mutations indicating highly parallel evolution or potential cross mixing). Notably, 5 different types of SNPs in *fusA*, encoding elongation factor G, were identified across all lineages and a *fusA* mutation was present in all samples sequenced (**Supplementary Data 1**). Additionally, 2 types of mutations in the upstream region of *abrB*, encoding transition state genes transcriptional regulator, were commonly mutated in 3 ALE experiments and their corresponding isolates (**Table 1**). Interestingly, one *abrB* mutation was a single SNP found 62 bp upstream of *abrG* (found in replicate lineage 5), and the other mutation identified was a combination of a 2 bp substitution (2 bp→TC) 52 bp upstream with a SNP 46 bp upstream (found in lineages 4 and 6). It is unclear if these mutation events happened at the same time during the experiment or if there was an order to their occurrence and this could be a subject of later analysis. Finally, there were single occurrences of intergenic SNP mutations in *walH*, a two-component regulatory system regulator of *WalRK*, in lineage 2 and a then a distinct SNP in *walK*, a cell wall metabolism sensor histidine kinase, in lineage 3. Both genes are in the same operon. Collectively, these convergent mutations suggest a strong influence of these genes on heat-shock tolerance^44^.

**Table 1.**
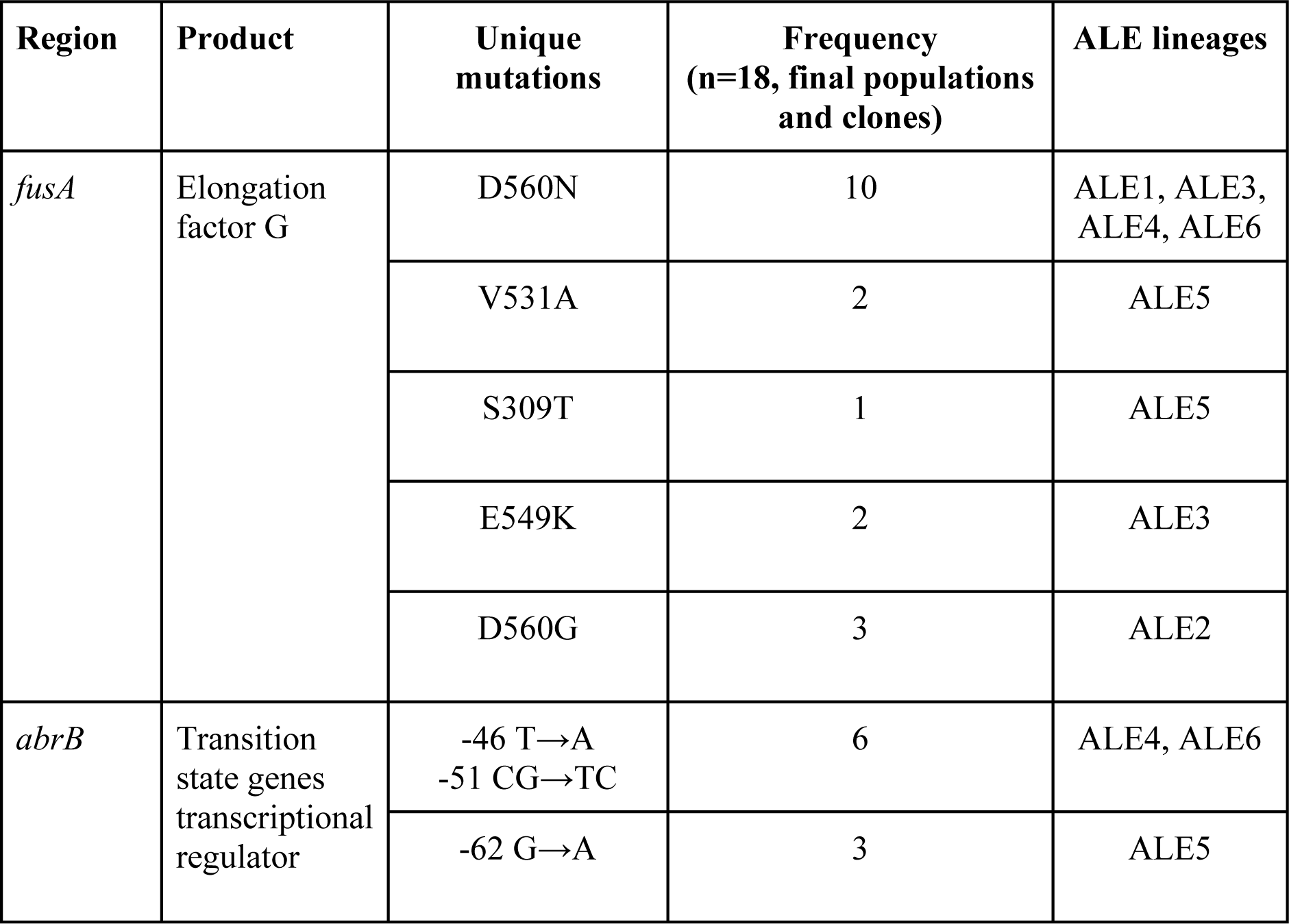
Commonly observed mutations from evolved isolates.

To directly assess the effects of mutation selected for related to heat tolerance, we used several isolates with single or double mutations of *fusA* and *abrB* genes (**Supplementary Data 1**). The *fusA* gene encodes a translation elongation factor that is involved in protein synthesis, while the *abrB* gene encodes a transcription regulator that is involved in the control of various cellular processes, including sporulation^45,46^ and stress responses^47^. Relative heat tolerances could be measured with germination efficiency after 10 minutes of heat treatment compared to wild-type ATCC 6633 strain (**Fig. 2D**). The isolate A3_F40_I1, with a single *fusA*^D560N^ mutation, the most frequently observed mutation, showed 3.0-fold increased germination efficiency compared to the wild-type strain. Meanwhile, isolate A4_F40_I1 had an additional *abrG* mutation along with *fusA*^D560N^, through which the effect of the *abrB* mutation could be indirectly confirmed. The additional *abrB* mutation could bring a modest increase in heat-shock tolerance to 4.5-fold. Similarly, several unique *fusA* mutation effects could be deciphered along with *abrB* mutation; A5_F40_I2 isolate with *fusA*^V531A^ and *abrB* mutations showed a 2.5-fold increase and A5_F40_I1 isolate with *fusA*^S309T^ showed significantly increased heat-shock tolerance to 17.7-fold. Further research is required to fully understand the mechanisms underlying the observed increase in heat-shock resistance in the evolved *Bacillus* spores, but the causality analysis shows that mutations in these two genes resulted in increased heat-shock tolerance.

### 2.3. Biocomposite TPU fabrication by using heat-shock tolerized spores

Wild-type (WT) and heat-shock tolerized (HST, A5_F40_I1 strain) ATCC 6633 spores were incorporated into TPUs as bioactive additives during HME^40,48^ (**Fig. 1**). Strain A5_F40_I1 was chosen for biocomposite fabrication as it showed the highest heat-tolerance among the isolates screened and it only possessed 2 distinct mutations in the commonly mutated *fusA* and *abrB* genes (**Supplementary Data 1**). TPU pellets and lyophilized spores, up to 1 w/w%, were fed into a twin-screw extruder and rigorously mixed at 135 °C at 36 rpm for 15 min, followed by extrusion through a slit die. As a result, biocomposite (BC) TPUs with WT or HST spores (BC TPU^WT^ or BC TPU^HST^) were fabricated (**Fig. 3A**). Spores possess an oblong shape with a ∼500 nm diameter and ∼1 µm length and, thus, can serve as a submicron TPU particulate filler if spores remain intact during the extrusion process (**Supplementary Fig. 2**). Both BC TPU^WT^ and BC TPU^HST^ displayed a light brown color following the incorporation of lyophilized spores (**Fig. 3A**). A uniform color distribution revealed proper mixing of spores and TPU melt during HME. It was spectrophotometrically determined that extracted spores from BC TPUs showed very consistent spectra to each other regardless of the sampling site from the extrudate, confirming uniform spore mixing (**Supplementary Fig. 3**).

**Fig. 3.**
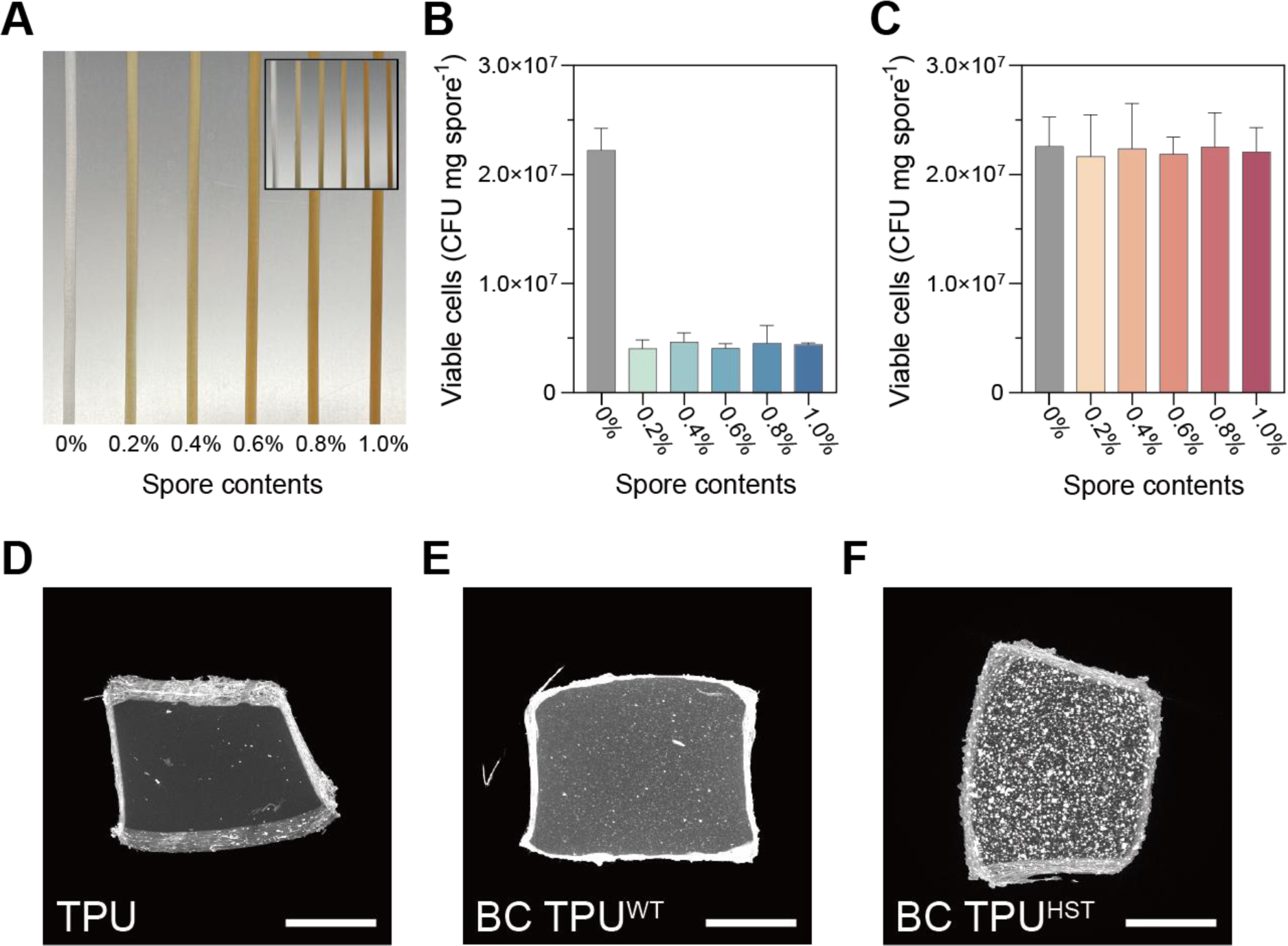
(A) Photographs of BC TPU^WT^ (inset) and BC TPU^HST^ with 0, 0.2, 0.4, 0.6, 0.8 and 1.0 w/w% spore loadings, left to right. The viability of WT (B) and HST (C) spores after HME. Spores were extracted from BC TPUs by dissolving the TPU compartment using DMF (no spore loss due to extraction assumed). Error bars indicate the standard deviations from three independent experiments. (D-F) MicroCT (XY projection) images of TPU and BC TPUs with 0.8 w/w% WT or HST spores (scale bars: 500 µm).

To quantitatively determine the viability of spores post HME, spores were extracted from the BC TPUs by dissolving the polymer component with DMF using an established protocol^49^. The viability of extracted spores was assessed by using a CFU assay. WT spores displayed ∼20% survivability after HME at 135 °C (**Fig. 3B**), whereas HST spores showed essentially full survivability (96-100%) regardless of spore loading (**Fig. 3C**). This finding indicated that the evolutionary engineering of *B. subtilis* spores by ALE markedly improved the heat tolerance of spores above the processing temperature of commercial thermoplastics, opening a new opportunity to fabricate biocomposite polymers. Full viability recovery after melt processing also suggested that the majority of the HST spores were highly resistant to the rigorous shear during HME (**Supplementary Fig. 4**). Notably, this enhanced heat tolerance was achieved without compromising any of the biological characteristics of the parental strain, as both of the WT and HST strains exhibited similar specific growth rates (∼0.8 h^−1^), spore yields (∼156 mg L^−1^), and specific cell viabilities (∼1.35 x 10^8^ CFU mg^−1^) when grown in pure liquid culture.

We visualized the 3-dimensional spore distribution in TPUs using micro-computed tomography (MicroCT) obtained using a X-ray microscope (XRM). XRM non-invasively revealed the successful incorporation of bacterial spores within the TPU matrices (**Fig. 3D-F & Supplementary Fig. 5**). Neat TPU has a high X-ray transmittance (∼80% transmission), hence the intensity of XRM is primarily attributed to the enriched calcium ions (Ca^2+^) in the spore core^50^. Interestingly, XRM images of BC TPU^WT^ vs. BC TPU^HST^ spores exhibited stark differences. BC TPU^HST^ showed clear contrast between the TPU background and bright particulate spores, while the X-ray signal of BC TPU^WT^ was relatively low and scattered across the matrix with smaller diameters compared to that of BC TPU^HST^. This finding was presumably due to structural damage of WT spores by heat and shear during HME. Ca^2+^ ions confined in the core of HST could be clearly detected, but XRM does not have sufficient resolution (700 nm spatial resolution and 70 nm voxel size) to detect dispersed Ca^2+^ ions from WT spore cores that were destroyed during processing. Well-defined particulates with high intensity in BC TPU^HST^ indicate a large population of HST spores retained their structure following melt processing.

### 2.4. Tensile properties of biocomposite TPUs

Spore incorporation generally had a positive effect on all mechanical properties measured for the BC TPUs. Tensile properties were evaluated based on four parameters; (i) toughness, (ii) elongation at break, (iii) ultimate tensile stress and (iv) Young’s modulus (**Fig. 4**) calculated from strain versus stress curves (**Supplementary Fig. 6**). For example, BC TPU^WT^ and BC TPU^HST^ showed up to 25% and 37% enhanced toughness, respectively, compared to TPUs without spores (**Fig. 4A & 4E**). This finding affirmed that the spores behaved as reinforcing fillers, improving the tensile properties of the TPU matrix. Toughness improvement of BC TPU^WT^ and BC TPU^HST^ was most pronounced at spore loadings of 0.4 w/w%, 0.6 w/w% and 0.8 w/w%, respectively. Additional loading above these critical concentrations led to a decreased toughness improvement, which was likely due to aggregation of spores within the matrix. High temperature and shear applied during the HME process, together with high spore concentration, has been found to induce the aggregation of spores in a TPU matrix^51^. Interestingly, the critical spore concentration of BC TPU^HST^ (0.8 w/w%) was higher than BC TPU^WT^ (0.4 ∼ 0.6 w/w%), while the ultimate toughness improvement of BC TPU^HST^ (37%) was also more remarkable than that of BC TPU^WT^ (25%) at their respective critical spore concentrations. These findings indicated that the heat tolerance of spores is important not only to retain their biological activities, but also for better filler behaviors.

**Fig. 4.**
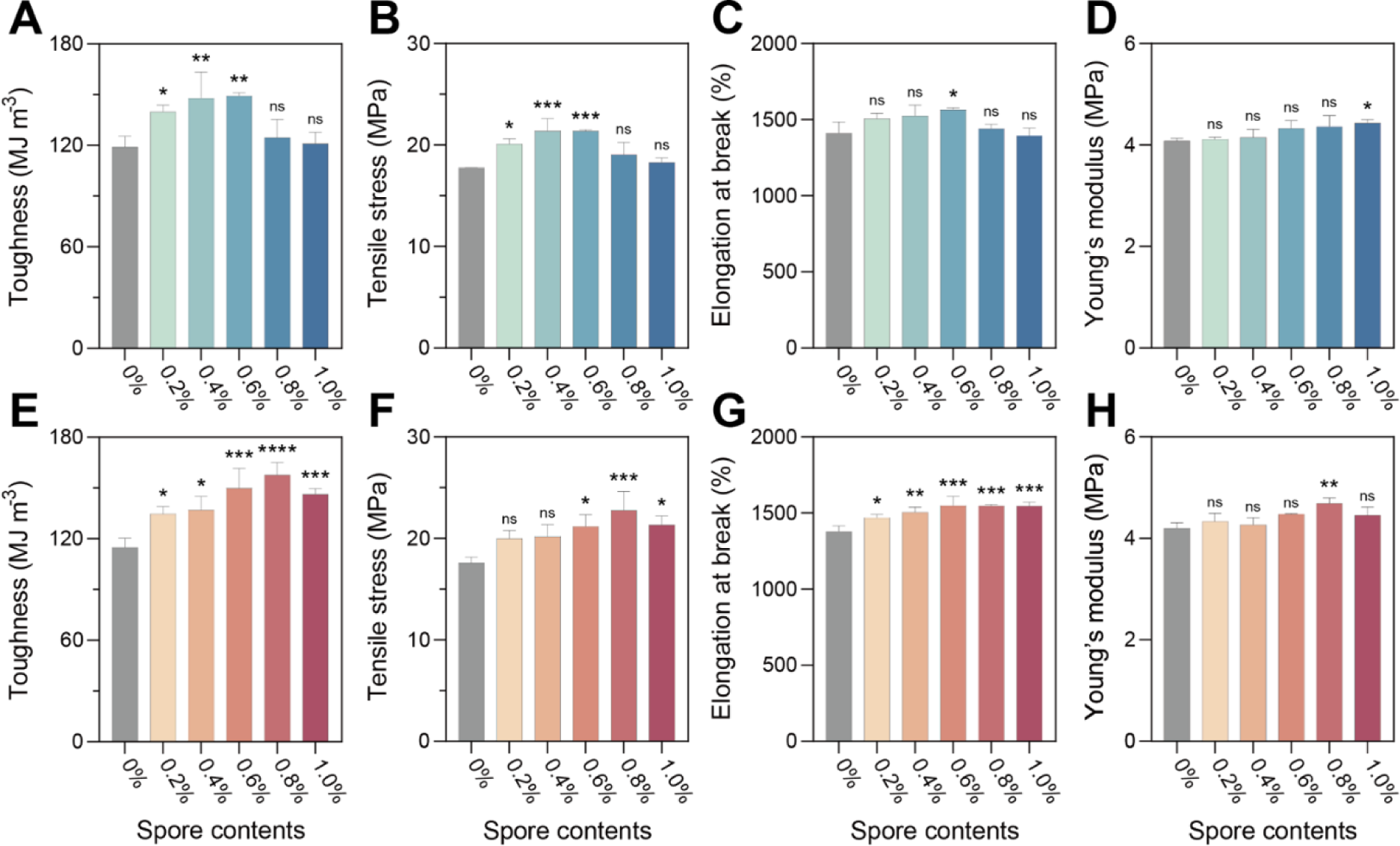
Tensile properties of BC TPU with WT (A-D) or HST spores (E-H). Error bars indicate the standard deviations from three independent experiments. One-way analysis of variance (ANOVA), followed by a post-hoc test with Dunnett’s multiple comparisons, was used for statistical comparison between TPU (control) and BC TPUs (n = 3 per group; ns: not significant; *P < 0.05; **P <0.01; ***P < 0.001; ****P < 0.0001).

Toughness improvements of spore-filled TPUs are primarily explained by the increase in tensile strength (**Fig. 4B & 4F**) and matrix yielding (**Fig. 4C & 4G**) of composite materials. Tensile stress and elongation at break of BC TPU^HST^ was increased by up to 30% and 12%, respectively, by spore addition (**Fig. 4F-G**). Similarly, BC TPU^WT^ showed up to 20% and 11% improvement in tensile stress and elongation at break, respectively, compared to TPU without spores (**Fig. 4B-C**). Increases in both the ultimate tensile stress and elongation at break of BC TPUs suggested a strong interfacial adhesion between the spores and TPU. Interaction between spores and TPU enhanced the energy dissipation in the composite systems based on the stress transfer, as well as prevented the interfacial cavitation during the viscoelastic deformation of composite materials^52^.

Interfacial adhesion between TPUs and spores was analyzed by an empirical model proposed by Pukanszky *et. al*^53^.

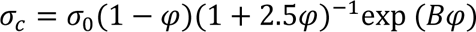

σ_c_ and σ_0_ are tensile stresses of filled and neat polymers, respectively. ѱ is a volume fraction of filler in the matrix, which was calculated from the weight fraction by using the densities of the TPU (1.18 g cm^−3^) and *B. subtilis* spores^54^ (1.52 g cm^−3^). B is the load-bearing capacity of the filler, which depends on polymer/filler interfacial interaction^55^. The Pukanszky model was rearranged as:

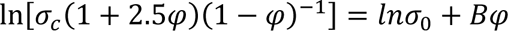

The B value was obtained from the linear correlation between ln[σ_c_(1+2.5ѱ)(1-ѱ)^−1^] and ѱ. Data before the critical spore concentration was used for the plot^52^. The B values of BC TPU^WT^ and BC TPU^HST^ were estimated as 23.6 and 31.4, respectively (**Supplementary Fig. 7**). Such high B values clearly depicted the strong polymer/filler interfacial adhesion^48^.

Even though both WT and HST spores served as reinforcing fillers for TPU with high B values, BC TPU^HST^ showed a higher B value compared to BC TPU^WT^, indicating stronger interfacial interaction between TPU and HST spores than that of TPU and WT spores. It can be explained by the (1) heat-shock tolerance of HST spores, which led to spores retaining their native structures (**Fig. 3D-F**) in TPU after HME and/or by (2) different interaction mechanisms between TPU and HST or WT spores. Water contact angle analysis revealed that the hydrophobicity of BC TPU^HST^ corresponded well with the toughness improvement profile along with the spore content (**Supplementary Fig. 8**). The finding implied that among potential TPU/spore interactions, the hydrophobic interaction could be a critical driving force of improved tensile properties, particularly for BC TPU^HST^. On the other hand, the hydrophobicity of BC TPU^WT^ remained similar regardless of the spore loading, suggesting that either the surface hydrophobicity of WT spores was less than that of HST spores or the denatured spores did not build a strong hydrophobic interaction with the TPU. This hypothesis is consistent with the different B values between BC TPU^WT^ (23.6) and BC TPU^HST^ (31.4), which implies that different types of interactions other than hydrophobic interactions may have contributed to the improved tensile properties of BC TPU^WT^. The Young’s modulus of BC TPUs was not significantly changed by the spore addition, suggesting that spores are soft filler materials and not affecting the stiffness of the TPU matrix (**Fig. 4D & 4H**).

### 2.5. Facilitated disintegration of biocomposite TPU

Facilitated disintegration of the BC TPUs was examined to determine the impact of spore incorporation, in general, and the impact that spore viability had post HME. This analysis is relevant as spores embedded in the BC TPU can remain dormant until germination is triggered, typically by the sensing of nutrients^20,56^, and reinforce the matrix as fillers throughout the lifespan of the TPU material. Then, at the end of the life-cycle of a BC TPU, spores can be germinated to facilitate TPU disintegration.

Approximately 80% of plastics are escaping our recycling efforts and being accumulated in landfills or the natural environment^57^. Furthermore, polyurethanes (PU) are the sixth most produced plastic in world^58^, but there is no governance for PU recycling. Even though PU waste can be potentially collected under the category seven of resin identification code (for miscellaneous plastics other than PETE, HDPE, PVC, LDPE, PP and PS), but only 0.3% of plastics in this category are generally recycled in the United States^59^. Furthermore, the global infrastructure for industrial composting in a microbe-rich environment is sparse and TPU waste would likely go to environments that are not enriched with TPU degrading microorganisms at the end of its lifecycle. Thus, it is important to test the degradation of TPU in microbially active and, more importantly, less active environments. We simulated these conditions by using microbially active compost and autoclaved compost, respectively. A CFU assay determined that untreated compost was enriched with viable cells (1.6 x 10^6^ CFU / mg compost), whereby a minimal number of microorganisms survived (0.0002%) following autoclaving (**Supplementary Table 1**). After confirming the successful germination and growth of spores using compost as a nutrient source (**Supplementary Fig. 9**), weight loss of BC TPUs in compost under controlled conditions of 37 °C and 45-55% relative humidity was assessed (**Fig. 5 & Supplementary Fig. 10**). BC TPU^HST^ exhibited significantly faster disintegration (92.7% mass loss) compared to TPU (43.9% mass loss) and BC TPU^WT^ (63.6% mass loss) in the autoclaved compost after 5 months (**Fig. 5A-B**). This suggested that both WT and HST spores could be germinated post HME and facilitate the disintegration of TPUs as living catalysts in an environment where few microbes were present when exposed to growth-supporting nutrients (**Supplementary Table 1**). Simultaneously, it indicated that the ∼100% survivability post HME in HST spores was strongly beneficial for the TPU disintegration kinetics in such conditions. Furthermore, BC TPU^HST^ in autoclaved compost showed comparable weight loss with that in unsterilized biologically active compost (**Fig. 5B & Supplementary Fig. 10B**). This finding indicated that the additional viability that HST provided can significantly aid in TPU fragmentation regardless of the inherent microbial activity in a given degradation environment. Disintegration of TPU control without spores in autoclaved compost can be explained by the re-growth of degrader strains in compost during the incubation and/or abiotic hydrolysis of TPU.

**Fig. 5.**
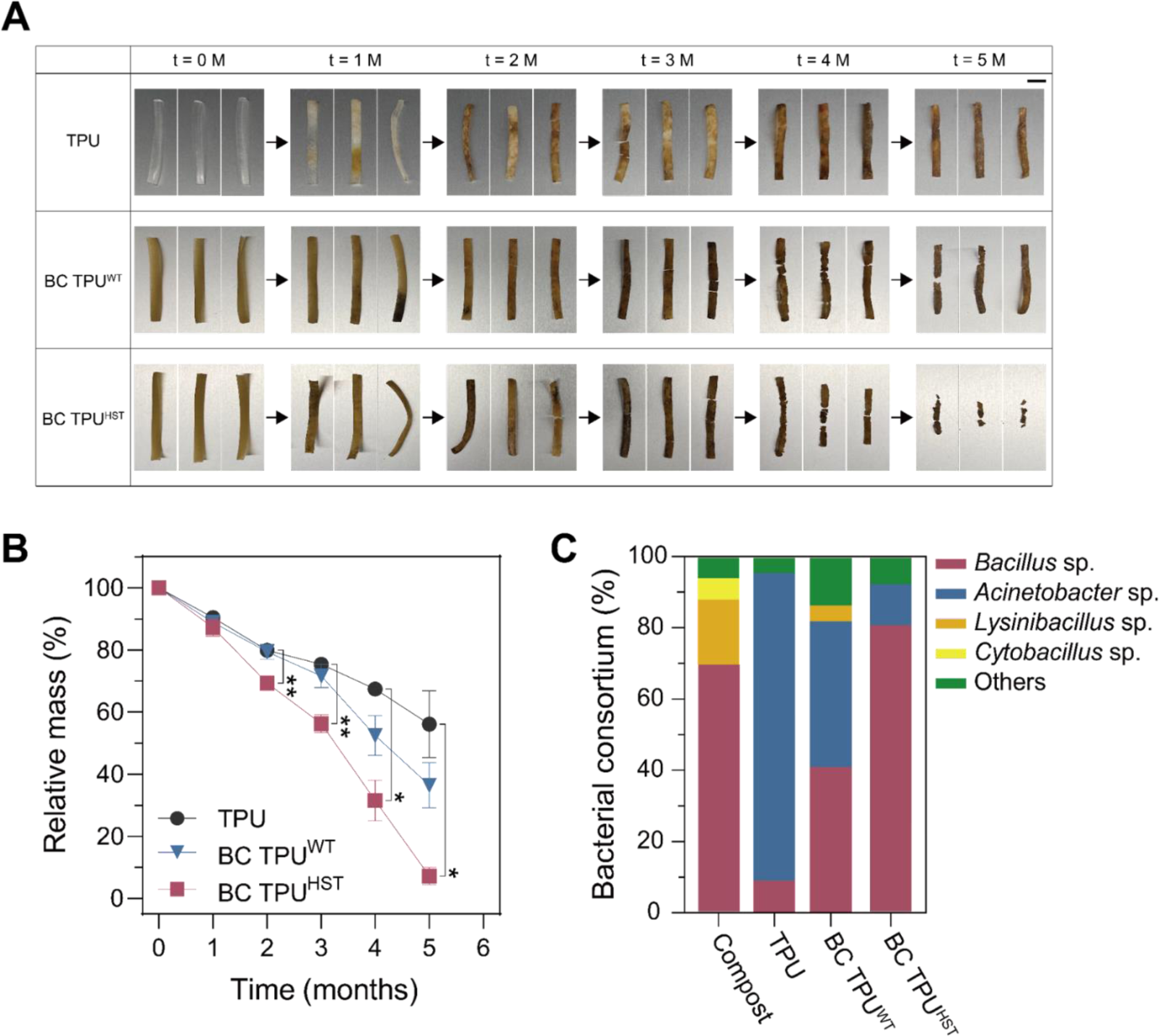
Photographs depicting the visual changes observed in the disintegration (A) and mass loss profile. (Scale bar: 1 cm) (B) of degraded TPU and BC TPUs during 5 months of incubation in autoclaved compost at 37 °C at 45-55% relative humidity (n = 3 per data point; *P < 0.05; **P < 0.01). Error bars indicate the standard deviations from three independent experiments. (C) Bacterial consortium analyzed from the compost and TPU surfaces incubated for 4 months in autoclaved compost.

To identify the microorganisms responsible for disintegration in compost, the surface of degraded TPUs was swabbed and analyzed using 16S rRNA sequencing (**Fig. 5C & Supplementary Fig. 10C**). It should be noted that the microbial analysis was focused on the bacterial community^60,61^ considering their abundance in compost and even after autoclaving, bacterial strains could be characterized in autoclaved compost, while the number of colonies were notably reduced (**Supplementary Table 1**). *Bacillus* sp, *Lysinibacillus* sp., and *Cytobacillus* sp. were commonly observed as dominant as over 90% in both compost samples (**Fig. 5C & Supplementary Fig. 10C**). Interestingly, *Acinetobacter* sp. was dominant on the surface of pristine TPU by 80.0% and 86.4%, respectively in untreated and autoclaved conditions after 4 months, which were below the limit of detection in the initial compost. This suggested that *Acinetobacter* sp. likely exhibited significant TPU disintegration activity among the microbial consortium^42,62^. Some *Acinetobacter* were observed in BC TPUs, however a clear increase in the population of *Bacillus* sp. was confirmed as expected. BC TPU^HST^ showed a higher proportion of *Bacillus* sp. (65.0% and 80.8% in untreated and autoclaved conditions, respectively) than BC TPU^WT^ (40.0% and 40.9%). Additional sequencing analysis revealed that 61.5% and 85.7% of the *Bacillus* sp. from BC TPU^HST^ were the evolved strain incorporated in HME. These results indicated that the enhanced viability of HST spores within the BC TPU contributes to their ability to germinate and colonize on the surface and participate in disintegration. Overall, these findings confirm that the HST spores can successfully germinate with biological activity after fabrication in compost and they indicate that the disintegration process is significantly accelerated in environments lacking sufficient quantity of degrader strains.

### 2.6. Programming biofunction into biocomposites through genetic engineering

Finally, the ability to genetically engineer ELMs is critically important to their overall ability to serve as smart materials. We genetically engineered the HST strain to demonstrate the introduction of a biofunction into biocomposite polymers. A plasmid harboring green fluorescent protein (GFP) was transformed into the HST strain and then incorporated into a BC TPU (BC TPU^HST/GFP^). Subsequently, BC TPU^HST/GFP^ were incubated in either PBS, LB, and compost extract (CE) along with control samples (TPU and BC TPU^HST^). The composite materials were then imaged using confocal laser scanning microscopy (CLSM) analysis.

CLSM imaging revealed the absence of green fluorescence signal from the TPU and BC TPU^HST^ incubated in PBS or LB (**Supplementary Fig. 12**). There were weak fluorescence signals observed from BC TPU^HST/GFP^ incubated in PBS (**Fig. 6**). This can be attributed to the residual GFP that was constitutively expressed during the bacteria cultivation and retained in or on spores during the spore purification process. In contrast, BC TPU^HST/GFP^ incubated in LB and CE exhibited strong GFP signals across the TPU matrix (**Fig. 6**). This indicated that the spores within the BC TPU successfully germinated and self-replicated, utilizing the available nutrients, leading to spontaneous GFP production. Furthermore, bright-field images obtained during CLSM analysis clearly showed the coincidence of spores and germinated cells on BC TPU^HST/GFP^ (**Fig. 6**), providing evidence of spore germination within the composite material triggered by nutrients. These results indicate the plasmid could be retained and protected along with chromosome within HST spores during the HME process, and the desired function can be harnessed with genetic engineering.

**Fig. 6.**
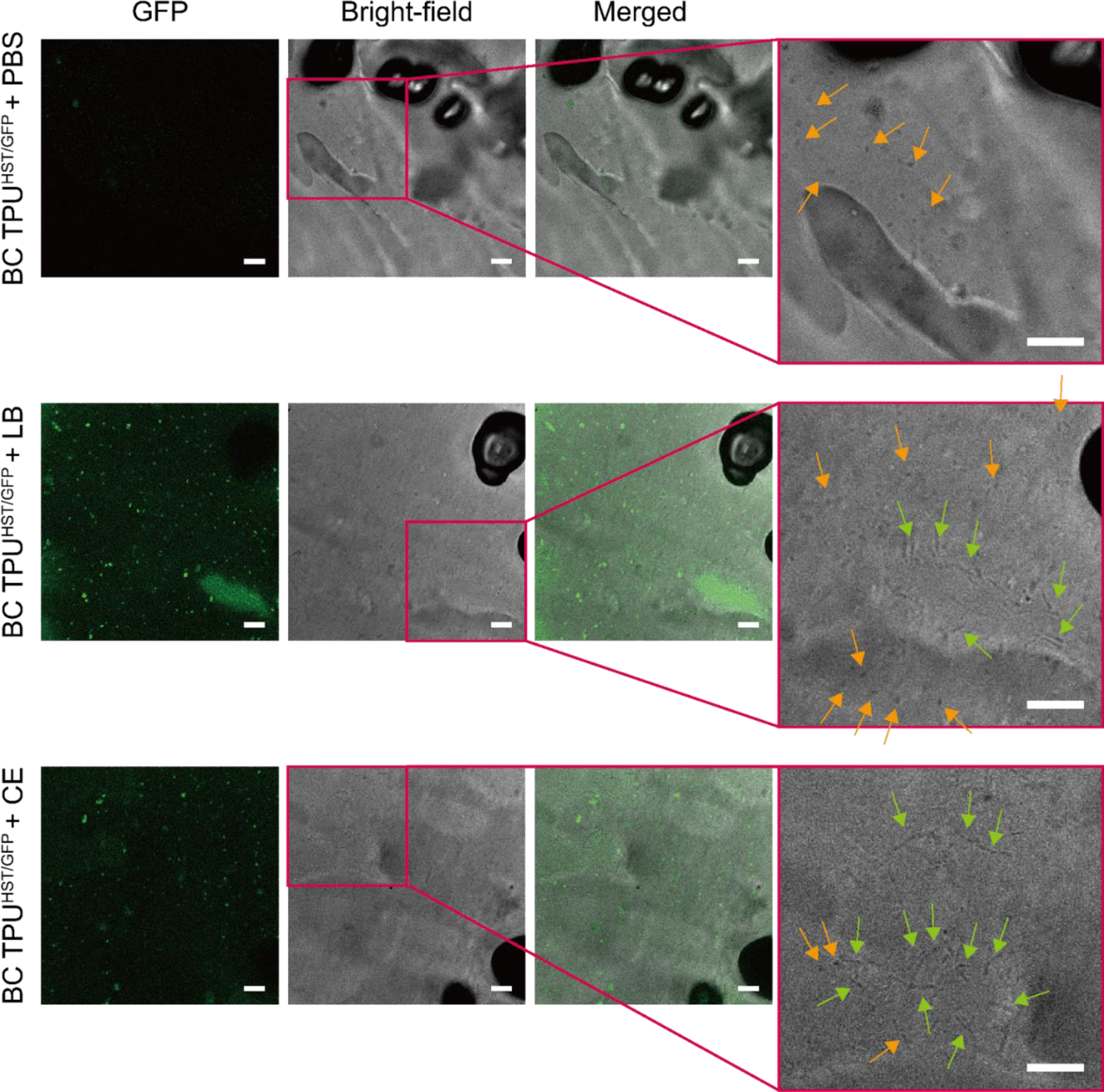
Fluorescence, bright-field and merged images (left to right) of BC TPU^HST/GFP^ incubated in PBS, LB, and CE obtained by CLSM. Rightmost panels are magnified bright-field images to identify the coincidence of rod-shaped vegetative cells with ∼ 5 µm length (green arrows) and particulate spores with ∼ 1 µm length (orange arrows). Scale bars: 10 µm.

## 3. Discussion

Engineered living materials hold great promise to expand the usability and applications of polymeric materials. However, there are fundamental cellular physiological constraints in ELM fabrication and use. Primarily, a number of polymer/cell interactions need to be overcome and understood to generate viable materials. In this work, evolutionary engineering was critical to generate functional BC TPU materials that were able to disintegrate and maintain a genetically engineered heterologous expression system. Accordingly, the main findings from this work were that, i.) the evolutionary engineering approach using a heat shock ALE approach was effective in generating spores that could survive the hot melt extrusion process and the genetic basis of these mutations could be determined, ii.) incorporation of heat-shock tolerized spores as living fillers into TPUs resulted in an overall increase in the tensile properties of TPUs owing to the strong interfacial interactions between spores and polymer, and, iii.) incorporation of spores into TPU to generate a living plastic material resulted in increased disintegration in compost. The results from each of these findings have implications for extension of the overall approach.

BC TPUs were fabricated by incorporating evolutionarily engineered *B. subtilis* spores into an HME process. ALE increased the heat-shock tolerance of *Bacillus* spores significantly without compromising overall fitness and strain usability (i.e., there were no apparent tradeoffs). Furthermore, evolved cells were shown to effectively express a heterologous protein which expands opportunities for future applications. Sequencing and causality analysis revealed that highly converged mutations (a surprising 44% of all unique mutations identified across six replicates) in *fusA* and *abrB* enhanced heat-shock tolerance. These findings are consistent with previous studies that have shown mutations in genes encoding translation^63^ and transcription factors^37,64,65^ can confer increased stress tolerance in bacteria. Such specific mutations and targeted genetic regions would have been extremely difficult to predict and engineer using rational approaches. A unique aspect of this approach is that evolutionarily-engineered microorganisms derived from ALE are regarded as non-GMO products given that ALE is facilitating and directing the natural evolution processes without artificial intervention^66^. These advantages suggest the feasibility of fabrication and application of industrial BC TPUs utilizing the presented strategy.

The incorporation of heat-shock tolerized spores as living fillers into TPUs resulted in an overall increase in the tensile properties of TPUs owing to the strong interfacial interactions between spores and polymer matrix. This enhancement in tensile properties of TPU by spore addition offers a promising solution to overcome the tradeoff barrier between tensile stress and elongation at break in commercial TPUs (**Supplementary Fig. 13**). The evolved bacterial spores exhibited essentially full viability even after the melt processing, clearly demonstrating a high level of biological functionality. The viability of spores following HME was critical for improvement of tensile properties and facilitating disintegration. Furthermore, we showcased the potential of genetic engineering by incorporating a GFP-expressing plasmid into the strains. It should be also noted that *B. subtilis* is not only beneficial for human health, but also aids the growth of plants as biocontrol agents^67^ and, thus, is considered a GRAS^68^ bioactive additive.

HME, used in this work, is an established technology for industrial polymer manufacturing and more than half of all plastic products are fabricated by HME^40,69^. The scale-up and scale-down of TSE-based HME is widely studied^70^, and the throughput of BC TPU fabrication (15 g/h) by the benchtop TSE (**Supplementary Fig. 14**) can be easily increased to larger scales. For example, our data showed that spores retain significant survivability (∼88 %) up to 144 rpm screw speed, which corresponds to up to 220 MPa shear stress (**Supplementary Fig. S15**). Increasing the screw speed, together with an increase in the screw geometry will achieve exponential upscaling. Under volumetric scaling, the throughput can be increased to the power of 3 with an increase in outer screw diameter (D)^71,72^.

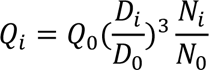

where, *Q* is throughput, *D* is outer diameter of screw and N is screw rotation speed. Subscripts i and 0 represent two different scales. In addition, BC TPU fabrication is aligned with the workflow of the current practice of industrial TPU manufacturing, as lyophilized spores are a compatible dry additive.

Incorporation of evolved spores into the tested TPUs resulted in an increased disintegration and informs how the approach can promote fragmentation at the end of lifecycle in an environment void of robust microbial actively, as would often be the case in consumer polymer disposal. Future steps to engineer the cells for enhanced disintegration should facilitate disintegration for the targeted TPU and likely others^73^. It was demonstrated that BC TPUs in compost with robust microbial activity did not show an advantage in disintegration, implying that this approach may not be necessary for such rich conditions. Nonetheless, the synergistic combination of genetic and evolutionary engineering can be further utilized to enhance TPU disintegration and introduce new biological functionalities to biocomposite materials.

Importantly, the practical use of TPUs relies on their mechanical properties. Thus, it is essential to maintain or improve these properties in developing biodegradable TPUs. The baseline TPU used in this work, BCF45, is a commercial grade TPU manufactured by BASF, which showed excellent tensile properties among TPU materials in the market (**Supplementary Fig. 13**). Spore incorporation not only facilitated the biodegradation, but also improved the mechanical properties of BCF45 (i.e. tensile strength ∼30%; elongation at break ∼20%; toughness ∼37%). There is a trade-off barrier between the tensile strength vs. elongation at break of commercial TPUs (**Supplementary Fig. 13**), however the incorporation of HST spores moved the mechanical properties beyond this apparent barrier. State-of-the-art biodegradable TPUs are generally designed by blending the TPU with biodegradable polymers, such as cellulose, polylactic acid or polycaprolactone, or by optimizing the raw components for TPU synthesis to incorporate bio-based feedstocks^74–79^. However, tensile properties of these materials have not surpased the trade-off-barrier^74–79^. Thus, BC TPUs developed in this work improve both tensile properties and biodegradation, which can potentially widen the application of TPUs.

The current manuscript focuses on addressing the fate of TPUs in nature. Autoclaved compost simulates soil or landfill with marginal biological activity, while untreated compost fosters microbially active condition for TPU biodegradation. Under both conditions, BC TPU^HST^ achieved over 90% biodegradation at 37 °C in 5 months regardless of the microbial activity of the degradation substrate. This result depicts that nutrient and moisture content were sufficient for facilitating biodegradation. In other words, spores embedded in the TPU matrix triggered and facilitated the biodegradation of TPU with minimal intervention. In comparison, most reports have tested the degradation of TPUs under specialized conditions^74–79^, which often do not reflect the realistic fate of polymer waste.

In conclusion, the incorporation of bacterial spores presents exciting opportunities for the introduction of living cells as renewable polymer fillers in industrial processes. This innovative approach combines evolutionary and genetic engineering methodologies and shows potential for diverse applications in the advancement of biocomposite materials.

## 4. Methods

### 4.1. Bacterial cells, plasmids, and materials

*Bacillus* strains were obtained from the American Type Culture Collection (ATCC, Manassas, VA, USA). Q5^®^ High-Fidelity DNA Polymerase, restriction enzymes, and NEBuilder® HiFi DNA Assembly Master Mix were purchased from New England Biolabs (NEB, Ipswich, MA, USA). Oligonucleotides for genetic manipulations were synthesized by Integrated DNA Technologies (IDT, Coralville, IA, USA). Soft-grade, polyester-based thermoplastic polyurethane pellets (Elastollan^®^ BCF 45) were gifted from BASF. All other chemicals were purchased from Sigma-Aldrich (St. Louis, MO, USA). To create GFP expressing plasmid, the Cas9 gene cassette in pAW016-2^80^ (Addgene, Watertown, MA, USA) was replaced with the GFP expression cassette. The plasmid was then introduced into HST strain^81^.

### 4.2. Cell culture for bacterial growth and sporulation

Luria–Bertani (LB, 10 g/L tryptone, 5 g/L yeast extract, 10 g/L NaCl) medium was utilized for routine cell culture. Dicfo Sporulation Media (DSM) containing 8 g/L Bacto nutrient broth, 1 g/L KCl, and 0.12 g/L was used to sporulate *Bacillus* strains. Freshly prepared metal ion solutions were added into DSM medium before use to the final concentration of 1 mM CaCl_2_, 1 µM FeSO_4_, and 1 µM MnCl_2_. To induce the germination of spores containing the GFP-expressing plasmid, erythromycin was included in the media at a final concentration of 5 μg/ml. This antibiotic was used for selection purposes to ensure the growth of spores that successfully incorporated the plasmid.

### 4.3. Adaptive Laboratory Evolution (ALE)

Cell cultures for ALE were performed with a 30 mL cylindrical tube containing 15 mL of DSM media at 37°C and stirred at 1100 rpm. Seed cultures were prepared by inoculating individual colonies from LB agar plates for each ALE experiment. Sporulations could be achieved with cultivation for 24 h in DSM media. Subsequently, the cultures were adjusted to a final concentration of OD_600_ 1.0 and subjected to heat-shock treatment by immersing the tube in boiling water (100 °C). Following the heat-shock treatment, the cell cultures were centrifuged at 5000 rpm for 10 minutes, and the resulting cell pellet was resuspended in fresh media for the subsequent passages, achieving a final concentration of OD_600_ 0.3 - 0.4.

### 4.4. Genome sequencing analysis

Genomic DNAs were prepared by using Mag-Bind^®^ Bacterial DNA purification kit from Omega Bio-Tek (Norcross, GA, USA) and converted into a sequencing library by using NEBNext^®^ Ultra™ II DNA Library Prep Kit (Ipswich, MA, USA). Sequencing was performed by using Illumina Novaseq 6000 (San Diego, CA, USA) at UC San Diego IGM Genomics Center. Sequencing raw reads were processed with Breseq (version 0.35.4)^82^ and a frequency cutoff ≥ 0.25 was applied for mutation analysis (**Supplementary Data 1**). The results were uploaded to ALEdb v1.0^83^ (http://aledb.org, project id: BS6633_HSTALE).

### 4.5. Spore production and purification

Glycerol stock of *B. subtilis* was inoculated into 25 mL of LB at 1.25% (v/v) to prepare seed culture. The seed culture was incubated at 37 °C at 200 rpm shaking overnight. 0.025L of seed culture was then mixed with 2.475 L of DSM for the main culture and subsequent sporulation. The inoculated DSM was evenly distributed into six baffled-flasks (4 L capacity) and incubated at 37 °C at 150 rpm shaking for two days in order to saturate and sporulate *B. subtilis*. *B. subtilis* spores were collected from the culture by the centrifugation at 8000 rpm at 4 °C for 20 min (Avanti J-E, Beckman Coulter Life Science, IN, USA). Supernatant was removed and the spore pellets were resuspended in fresh 100 mM PBS pH 7.4. The recovered spores were washed by repeating the following steps for three times: (i) centrifugation at 4000 rpm at 25 °C for 10 min (5810 R, Eppendorf, Hamburg, Germany), (ii) removal of supernatant, and (iii) resuspension of spore pellet in fresh 100 mM PBS pH 7.4. The spores were then treated with 2.5 mg/mL lysozyme solution at 37 °C at 150 rpm shaking for 1 h, followed by three times of washing with 100 mM PBS pH 7.4. The spores were further heat-treated at 65 °C for 1 h under static condition. After three times of washing by using deionized water, the spores were suspended in deionized water, frozen by using liquid nitrogen for 5 min, and thoroughly lyophilized for two days (FreeZone 2.5, Labconco, MO, USA).

### 4.6. Scanning electron microscopy (SEM)

SEM analysis of lyophilized spores were conducted by using FEI Apreo 2 SEM (Thermo Fisher Scientific, Waltham, MA,USA). Spores were coated with gold for 60 sec for 3 times. Images were obtained under high vacuum at 20 kV at 3.2 nA.

### 4.7. Hot melt extrusion (HME)

Biocomposite TPUs were fabricated by using a twin screw extruder (HAAKE^TM^ MiniCTW, Thermo Fisher Scientific, Waltham, MA, USA) equipt with a slit die. ∼ 5 g of TPU pellets were loaded to the twin screw extruder and incubated at 135 °C at 36 rpm for 5 min under cycle mode. Then, the lyophilized spore powder was introduced to the twin screw extruder at various loading ranging from 0 to 1.0 w/w%. The TPU-spore mixture was further incubated at 135 °C at 36 rpm for 15 min under cycle mode, and then extruded at 3 rpm under flush mode.

### 4.8. UV-Vis spectrophotometry

Three 100 mg pieces of biocomposite TPU with 0.8 w/w% spore loading were collected from different sites in an extrudate. Each biocomposite TPU aliquot was dissolved in DMF. UV-Vis absorbance spectra of dissolved biocomposite TPUs and other controls were obtained at 260 ∼ 700 nm by using a microplate reader (Synergy HT, BioTek, Winookski, VT, USA)

### 4.9. X-ray microscopy (XRM)

MicroCT images of TPU and biocomposite TPUs were obtained using Xradia 510 Versaat (Zeiss, Oberkochen, Germany) operated at 80 kV and 7 mA. All the samples were analyzed without staining. Neat TPU and biocomposite TPUs were visualized at 4x magnification. X-ray exposure time was 1 second. 3D projection images and 3D volumes of microCT were constructed from tilt series with 2401 projections using XMController and XMReconstructor, respectively (Xradia).

### 4.10. Tensile testing

Biocomposite TPU was tailored into a dogbone shape by using a guided cutting method for the tensile testing (overall length: > 38 mm; clamping area length: > 10 mm; initial distance between grips: ∼18 mm; length of narrow parallel-sided portion: 10 mm; width at ends: 5 mm; width at narrow portion: 2.4 mm; thickness: 0.7 mm). The dogbone specimen was loaded between two grips with serrated jaws. The specimen was stretched at 20 mm/min extension rate until it fractured by using a universal testing machine (Instron 5982, MA, USA). Load (N) vs. extension (mm) curves obtained by the tensile testing were converted into stress (MPa) vs. strain (-) curves using the following equations.

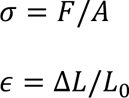

where, σ is stress, F is load, A is cross-sectional area of the specimen, ɛ is strain, ΔL is displacement and L_0_ is the initial length of the specimen.

Ultimate tensile stress and elongation at break of the specimen were obtained from the maximum stress value and strain value at the fracture moment, respectively. Toughness was calculated from the area under stress vs. strain curve based on the trapezoid rule. Young’s modulus was calculated from the slope of stress vs. strain curve during the initial stretching of the specimen.

### 4.11. Shear stress calculation

Shear stress during the melt processing was calculated from the following equations.

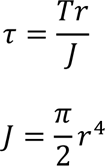

where, τ is shear stress, T is applied torque (obtained from MiniCTW), r is radius of shaft, J is polar moment of inertia.

Since conical screws were used for HME, radial distance (r) varied along with the length of screw. Thus, the maximum and minimum shear stresses were separately calculated from the minimum and maximum screw radii, respectively.

### 4.12. Water contact angle

Prior to the water contact angle analysis, TPU samples were flattened by using a hot press. A small piece of TPU sample (∼ 10 x 10 x 1 mm^3^) was wrapped in aluminum foil and pressed between two flat heating blocks at 120 °C for 10 min through pneumatic pressure. Flattened TPU sample was transferred to a glass slide. The glass slide was loaded to a contact angle goniometer (Rame-Hart 500, NJ, USA). A droplet of water was dispensed and the contact angle was measured from the image taken by the contact angle goniometer.

### 4.13. Spore extraction from biocomposite TPU

100 mg of biocomposite TPU was diced into small pieces (∼ 1 x 1 x 1 mm^3^), and incubated in 10 mL DMF at 40 °C under 150 rpm magnet stirring for 1 h to dissolve polymer compartment. The solution was then centrifuged at 6000 rpm at 25 °C for 10 min. Supernatant was removed and recovered spore pellet was resuspended in 10 mL of fresh DMF. The spore suspension was incubated at room temperature at 50 rpm rocking for 10 min. After removing DMF by the centrifugation at 6000 rpm at 25 °C for 10 min, the spore was dispersed in 100 mM PBS pH 7.4.

### 4.14. Cell viability test

The viability of spores was determined by colony forming unit (CFU) assay on LB plates. The viability could be accessed with 100 µL of serially-diluted spore suspension in 100 mM PBS pH 7.4 was spreaded onto LB plates. After 12 h incubation at 37 °C, the number of colonies appearing on LB plates was counted either manually or with image analyzing software (ImageJ). The concentration of viable cells in original spore suspension was calculated by considering the dilution factor.

### 4.15. Spore germination on compost gel

Compost was thoroughly dried at 80 °C under vacuum overnight. Dried compost was sieved by using a 35 standard mesh screen to yield fine powder. Dried compost and agarose were suspended in deionized water, followed by the autoclave at 121 °C for 20 min. The final concentration of compost powder in suspension was varied from 0 g/L to 100 g/L, while the concentration of agarose was fixed at 15 g/L. After the autoclave, the liquid part of the mixture was gently taken by using an electrical pipette and transferred to petri dishes. Each petri dish contained 15 mL of compost extract-agarose mixture solution. The mixture was gelated at room temperature. Compost gel plates were flipped upside down and stored at 4 °C until use. Germination of spores on compost gels followed the same procedure with that on LB plates.

### 4.16. Gravimetric disintegration test

Raw compost aged 4 – 5 months was collected from two industrial composting facilities located in Athens, GA. Landscaping and forest residues, food waste, and livestock manure are inputs for both facilities. The temperature of the composting pile at sampling depth (approximately 30 – 100 cm) was 46 ± 5 °C. The compost was particle sieved through a 4.76 mm screen. The compost was mixed thoroughly with a resulting pH of 7.4 at the beginning of testing. See **Supplementary Table 2** for compost details.

Initial mass of each TPU piece used for the gravimetric disintegration test was 200 mg. At least 30 pieces of each TPU sample were incubated in the untreated and autoclaved compost. Autoclaved compost was prepared by autoclaving compost at 121 °C for 20 min. 9 pieces of TPU samples were incubated in 500 g compost (by dry weight basis) in each 2 L plastic container, which was drilled with ¼ inch air holes. The plastic containers were then placed in an incubator at 37 °C at 45∼55% relative humidity. The interior of the incubator was regularly disinfected using 70% ethanol after removing the plastic containers to prevent the (cross)contamination of autoclaved compost. 3 pieces of each TPU sample were collected every month. Large fragments of TPU were carefully collected from the compost, and the small fragments potentially remained in the compost was recovered after thoroughly drying compost, followed by sieving with a 35 standard mesh screen. The collected TPU samples were excessively washed with deionized water and dried at 80 °C under vacuum for overnight. The weight of degraded TPU samples was measured.

### 4.17. Respirometry evaluation of TPU degradation

Respirometry evaluation was carried out using respirometry (ECHO, Slovenske Konjice, Slovenia) to evaluate the extent of mineralization of the TPU. (see **Supplementary Table 2** for compost details). The evolution of CO_2_ was quantified to determine the extent and rate to which sample carbon is consumed by microbes in the aerobic composting environment. Industrial composting conditions outlined in ASTM D5338 – 15/ISO 14855 maintain temperatures of 58 ± 2 °C for the duration of the testing; however, the conditions were modified to 42 ± 0.5 °C to study the metabolism of mesophilic microbes. Moreover, industrial composting temperatures often fall below 42 °C during the maturation phase. Respirometry was carried out using untreated compost only because there is no reliable method to suppress the contamination of autoclaved compost in respirometer under the current practice. 7 g of each TPU, BC TPU^WT^ and BC TPU^HST^ was cryo-milled and mixed with 125 g of compost (by dry weight basis). Bioreactors were purged continuously with 200 mL/min air held at 42 ± 0.5 °C and stirred once weekly to avoid channeling and maintain moisture at 45 – 60 %. Methane concentrations were measured and found to be negligible. The evolved CO_2_ for each sample, *s*, was calculated daily by the difference in CO_2_ production (mg/day) of each reactor, *r,* compared to the daily average CO_2_ contribution from blank controls, *b*, as described by the following equations:

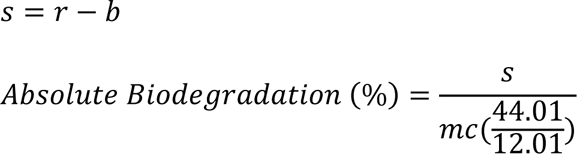

The sample mass, *m*, and the percent organic carbon content was used to determine the carbon content, *c*. The organic carbon and nitrogen contents of the polymers were determined with a Vario Elementar EL/max elemental analyzer using a TCD detector (Elementar Analysensysteme GmbH, Hanau, Germany) (**Supplementary Table 3**). The dimensionless value of 44.01 / 12.01 is used to account for the organic carbon in the CO_2_ generated from each reactor. Relative biodegradation was calculated from absolute biodegradation of samples relative to that of cellulose, a positive control.

### 4.18. Microbial consortium analysis in compost and composting experiments

To analyze the bacterial consortium, samples of compost or TPU obtained from the composting environment were collected. Sterilized swabs were used to gently swipe the samples onto LB agar plates. The plates were then incubated at 37°C overnight. From each sample, at least 20 individual colonies were randomly selected. These colonies were subjected to characterization using 16S rRNA sequencing (Eton Bioscience, San Diego, CA, USA/ Macrogen, Seoul, South Korea) with primers (27F - AGAGTTTGATCMTGGCTCAG, 1492R-TACGGYTACCTTGTTACGACTT). The obtained sequences were compared to existing databases using BLAST (blast.ncbi.nlm.nih.gov) to identify the bacterial species. For the clones identified as *B. subtilis*, additional sequencing was performed using primers (fusA_F - ATGGCAAGAGAGTTCTCCTTAGACAAAACT, fusA_R - GATAATTTCTTCTGAAACGCTCTTCGGCAC) to ensure the embedded strains.

### 4.19. Compost extract preparation

Compost extract was prepared by suspending dried compost powder in PBS at 100 g/L final concentration. The compost suspension was autoclaved at 121 °C for 20 min, followed by the sedimentation under static condition for 30 min. The liquid part was carefully aliquoted for further experiment.

### 4.20. Confocal laser scanning microscopy (CLSM)

Before CLSM analysis, TPU, BC TPU^HST^ and BC TPU^HST/GFP^ were diced into small pieces (∼ 1 x 1 x 1 mm^3^) and incubated in PBS, LB or compost extract at 37 °C under 150 rpm shaking overnight. 5 µg/L erythromycin was added to the media for BC TPU^HST/GFP^. The samples were then briefly rinsed with sterile PBS, then visualized using Leica SP8 confocal microscope equipped with white light laser (Leica Microsystems) at 100x magnification. The channel for GFP detection was set for excitation at 484 nm and the emission was collected at 497-594 nm. Additional bright-field images were recorded. Multiple images from independent regions of each sample were recorded. CLSM images were post-processed using ImageJ. GFP signals of all images were enhanced by uniformly adjusting brightness/contrast threshold at 0∼100. Minimum values of brightness/contrast threshold for bright-field images were fixed at 50 and the maximum values varied from 100 to 150 depending on the samples.

### 4.21. Statistical analysis

Analysis of variance (ANOVA), followed by a post-hoc test with Dunnett’s multiple comparisons, was used for the statistical analysis. For the tensile properties, one-way ANOVA was carried out and the mean of each group was compared with a control group (TPU without spores). Two-way ANOVA was performed for the disintegration data with a factor of time and sample. GraphPad Prism 9.2 was used for the statistical analysis. *P* < 0.05 was considered statistically significant, and additional indicators of statistical significance are provided accordingly in the text or in individual figure legends.

## Data Availability

The raw resequencing data generated during this study is available through the European Nucleotide Archive and National Center for Biotechnology Information Sequence Read Archive (BioProject number: PRJNA981571). The variant calling data is available at https://aledb.org (project name: BS6633_HSTALE). Other data that support the findings of this study are available from the corresponding authors upon reasonable request.

## Acknowledgements

This work was primarily sponsored by funding from BOTTLE^TM^ consortium (# DE-EE0009296) supported by the U.S. Department of Energy’s (DOE’s) Office of Energy Efficiency and Renewable Energy (EERE) and Advanced Manufacturing Office (AMO). This work was also in part sponsored by UC San Diego Materials Research Science and Engineering Center (UCSD MRSEC) (# DMR-2011924). The authors acknowledge the use of facilities and instrumentation supported by UCSD MRSEC (# DMR-2011924), UCSD NanoEngineering Materials Research Center (NE-MRC), National Center for Microscopy and Imaging Research (NCMIR) and UCSD School of Medicine Microscopy Core (# NINDS P30NS047101). We also acknowledge the XRM expertise and operation consulting offered by Dr. Guy Perkins and Dr. Keun-Young (Christine) Kim from NCMIR.

## Author Contributions

H.S.K. and M.H.N. designed and performed experiments and analyzed the results. M.A.R. conceived the project and selected TPU material. E.W., A.W., M.K., and J.L. prepared compost, performed respirometry analyses, and consulted on gravimetric disintegration tests. D.D. performed CLSM. The overall project was supervised by A.F. and J.P.. H.S.K., M.H.N., A.F. and J.P. wrote the manuscript with input from all authors. All authors contributed to the discussion of the results and the text.

## Competing Interests

All authors declare no competing interests.

## Notes

### Competing Interest Statement

The authors have declared no competing interest.

