## Supplementary Material for "Biocomposite Thermoplastic Polyurethanes Containing Evolved Bacterial Spores as Living Fillers to Facilitate Polymer Disintegration"

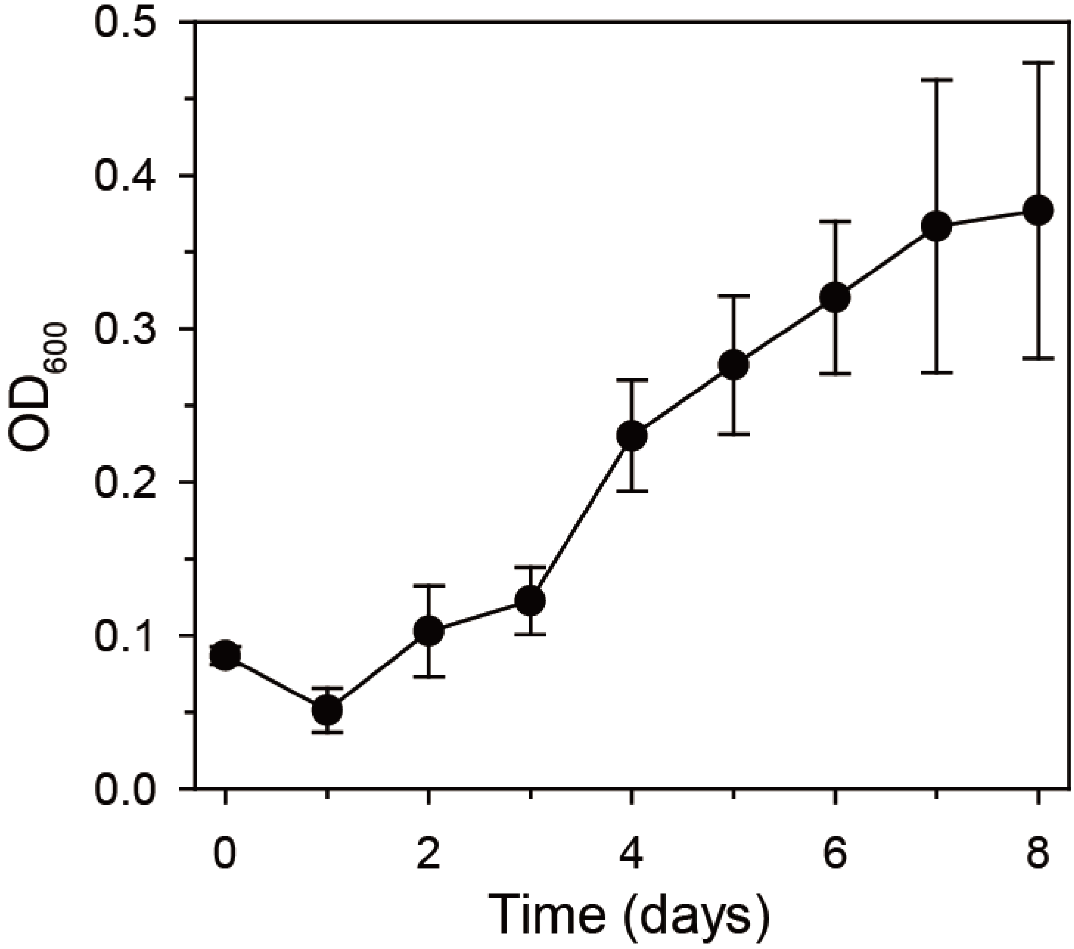

**Supplementary Fig. 1.** Growth of *B. subtilis* ATCC 6633 strain ($OD_{600}$) by the assimilation of polyester-based TPU as a sole carbon source. Error bars indicate standard deviations from three independent experiments.

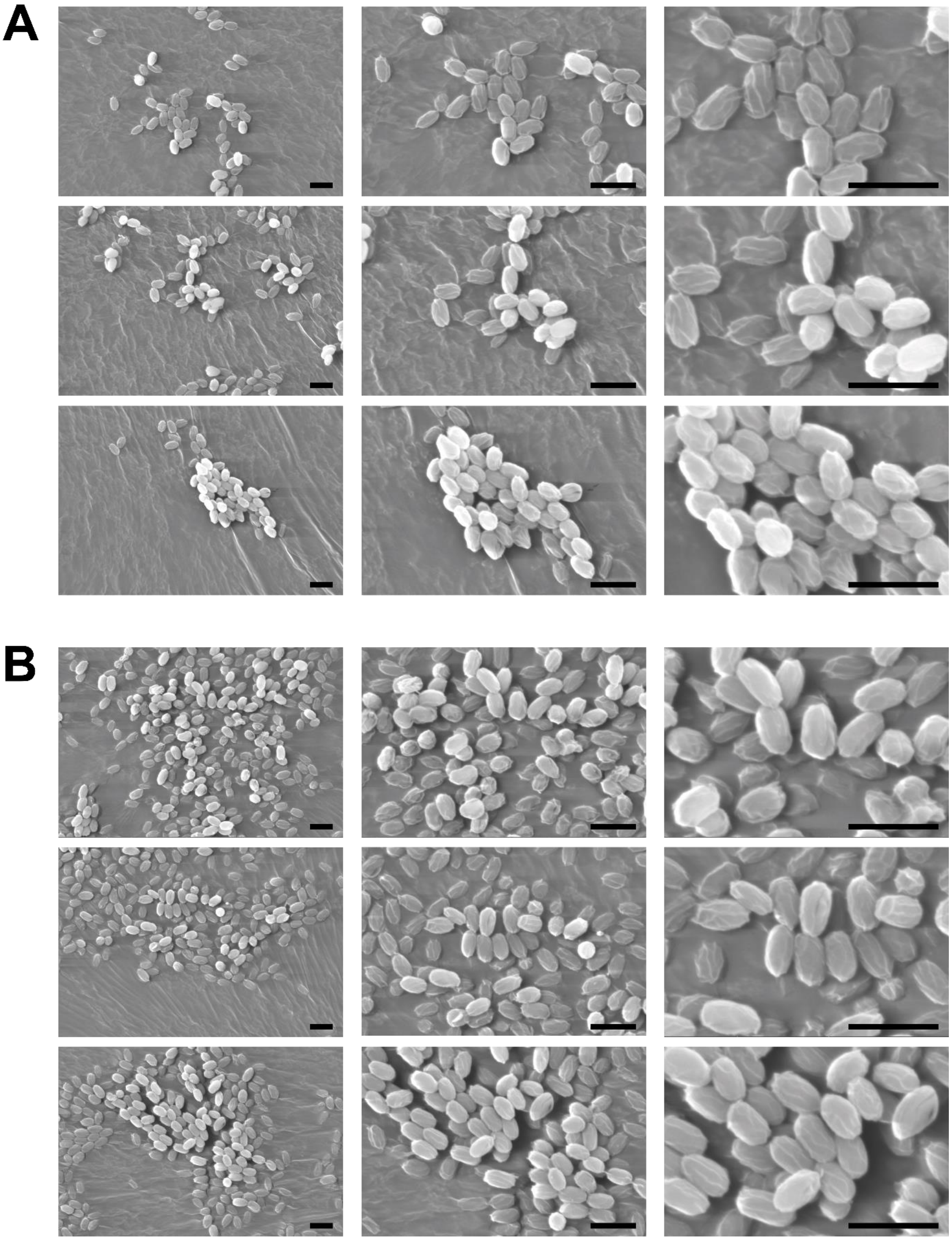

**Supplementary Fig. 2.** Scanning electron microscope images of lyophilized spores obtained from three different sites. (A) WT and (B) HST spores. Scale bars are 2 µm.

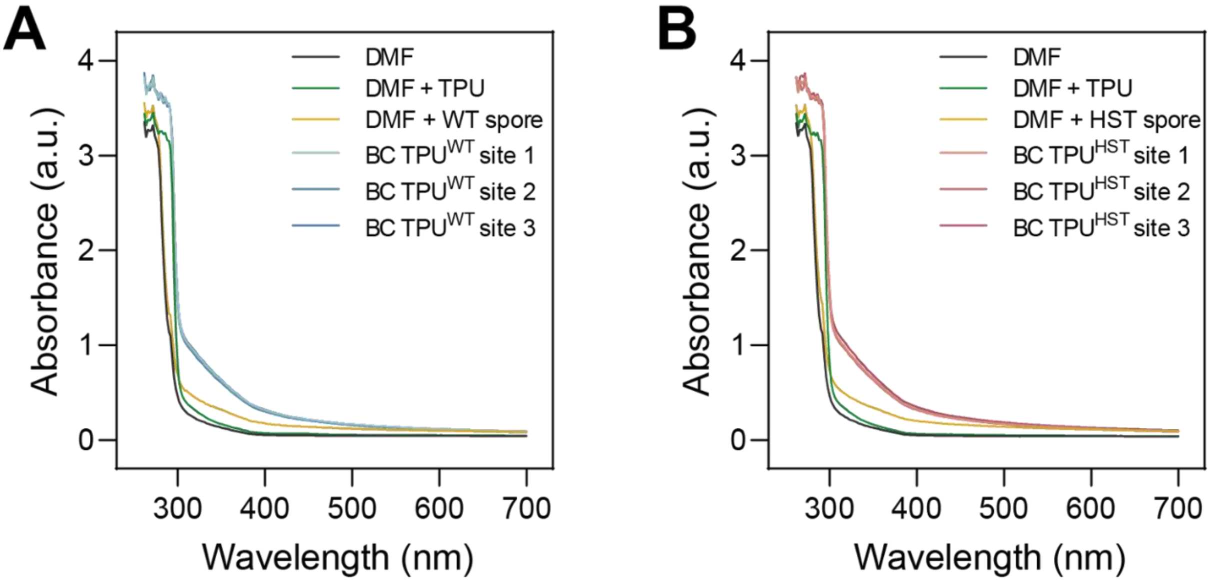

**Supplementary Fig. 3.** UV-Vis spectra of TPU, spores and BC TPUs with 0.8 w/w% spore loading dissolved/suspended in DMF. (A) WT and (B) HST. Final concentrations of TPU and spore in DMF were 9.92 mg/mL and 0.08 mg/mL, respectively. TPU showed absorbance at ~ 300 nm, while spores absorbed a broad range of UV-Vis after 300 nm. BC TPUs showed characteristic absorbance patterns from both TPU and spore. BC TPUs samples were collected from three different sites of an extrudate to confirm the uniform mixing of TPU and spores.

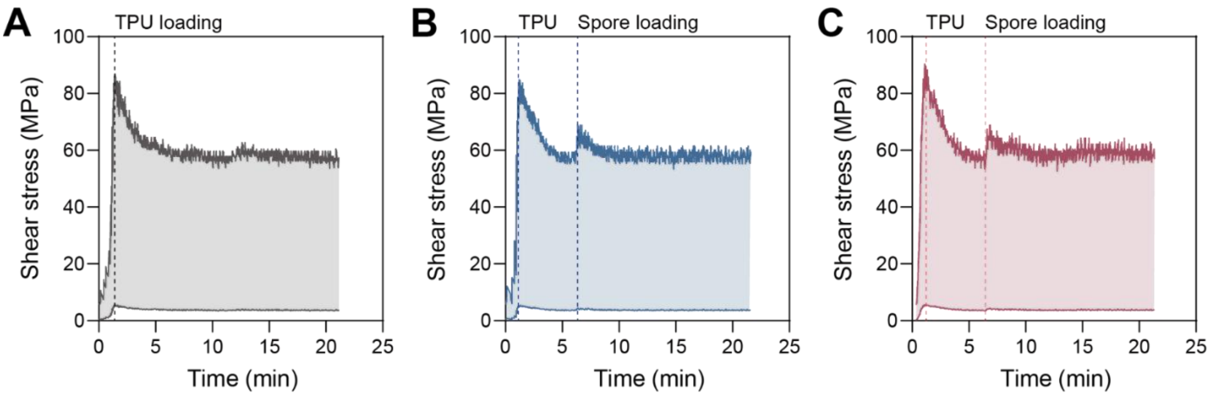

**Supplementary Fig. 4.** Shear stress during melt processing of TPU (A), BC TPU$^{WT}$ (B) and BC TPU$^{HST}$ (C). Spore loadings of biocomposite TPUs were 0.8 w/w%. Spores were added after 5 min of TPU melting and equilibration in the twin screw extruder for biocomposite TPU fabrication.

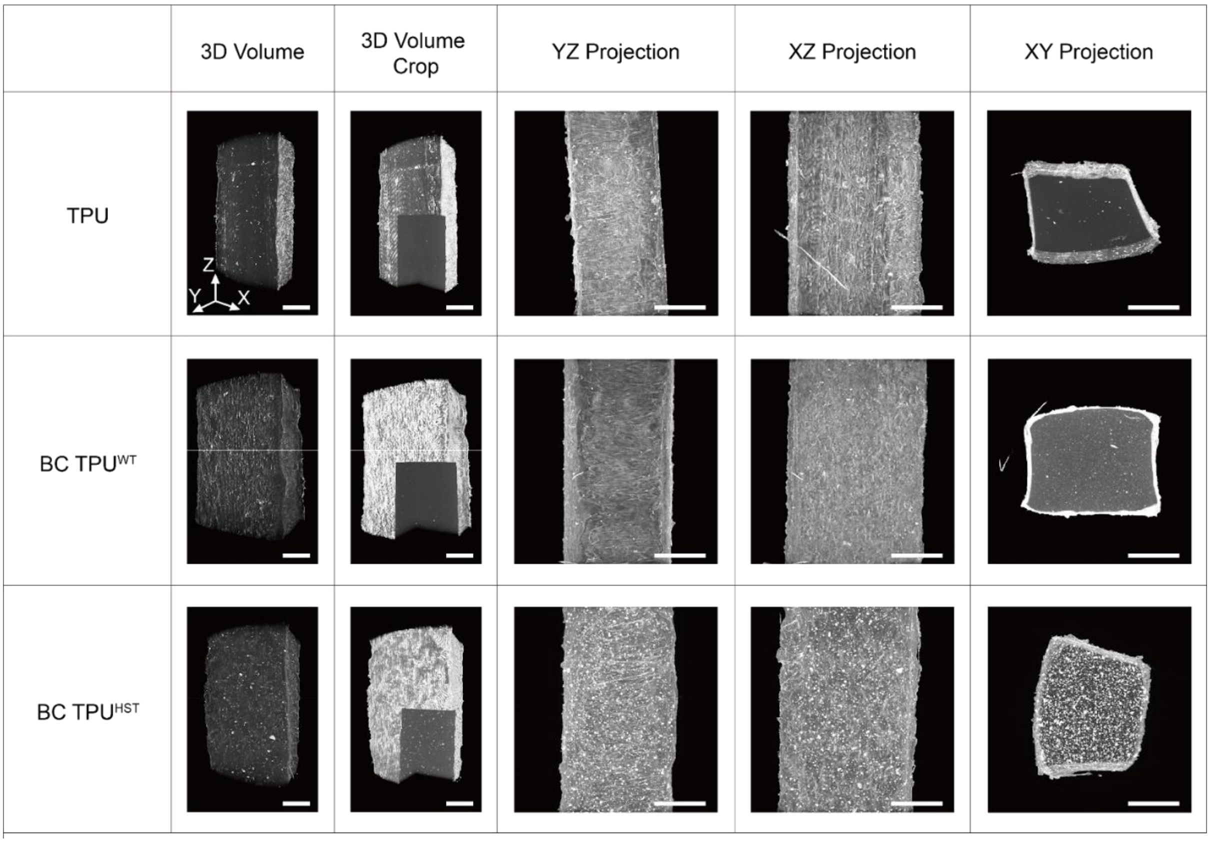

**Supplementary Fig. 5.** MicroCT images of TPU, BC $TPU^{WT}$ and BC $TPU^{HST}$ were obtained using XRM (scale bars: 500 µm). Spore loadings of biocomposite TPUs were 0.8 w/w%. Specimens for XRM analysis prepared at ~ 1 x 1 x 10 $mm^3$ (X x Y x Z) dimension. 3D scanning was conducted by rotating the specimens in the Z axis. Since X and Y dimensions of each specimen were smaller than the scanning frame, the rough surface of the specimen was visualized in MicroCT images except XY projection.

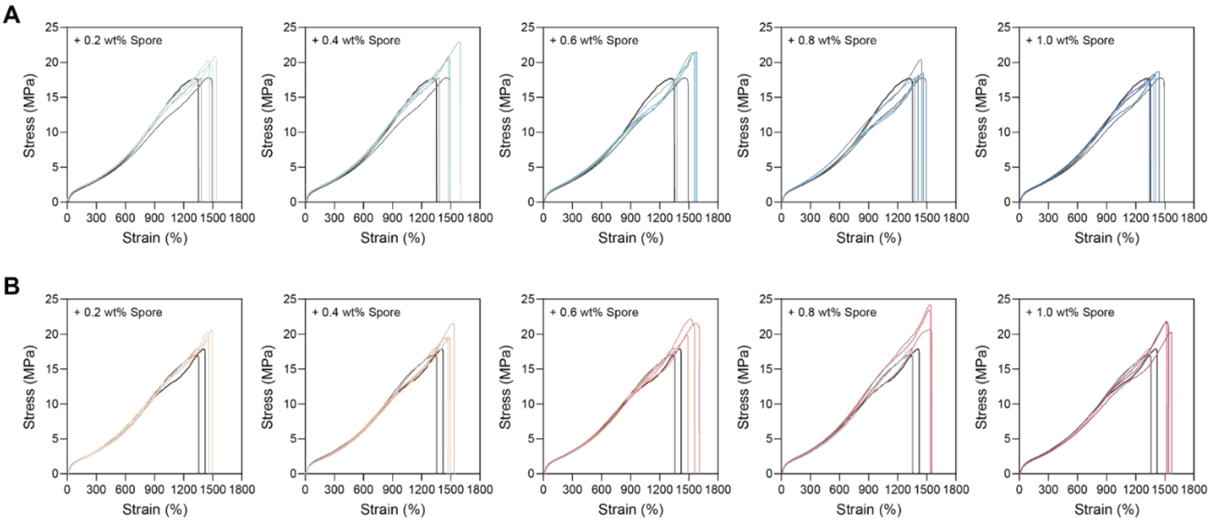

**Supplementary Fig. 6.** Stress versus strain curves obtained by the tensile testing of BC TPU$^{WT}$ (A) and BC TPU$^{HST}$ (B) with varying spore contents. Grayscale and colored curves represent TPU and biocomposite TPUs, respectively. Toughness was calculated from the area under the curve. Ultimate tensile stress was obtained by the Y peak, while the elongation at break was the strain at the moment of fracture. Young’s modulus was calculated from the slope in the initial range.

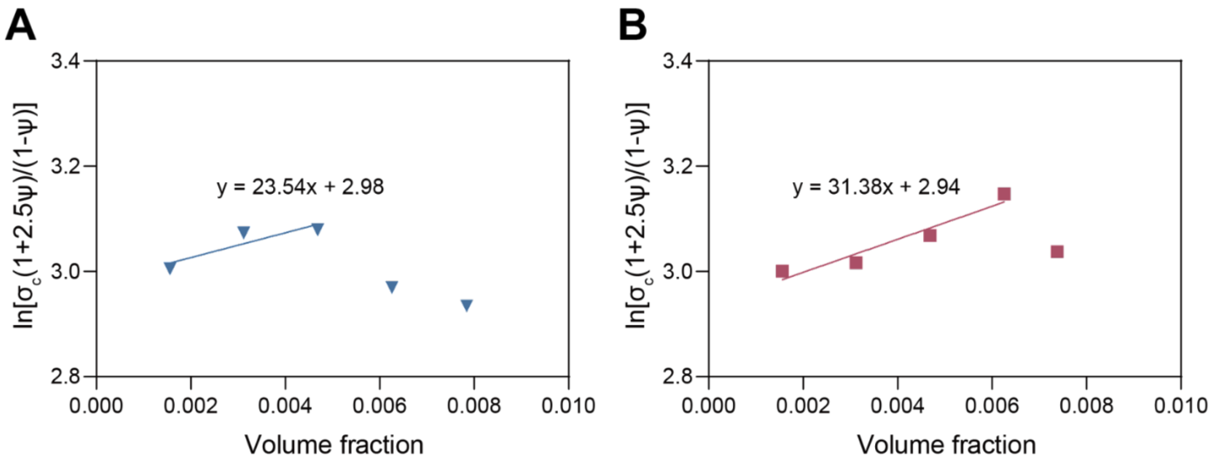

**Supplementary Fig. 7.** B values of BC TPU$^{WT}$ (A) and BC TPU$^{HST}$ (B) were calculated based on the Pukánszky model.

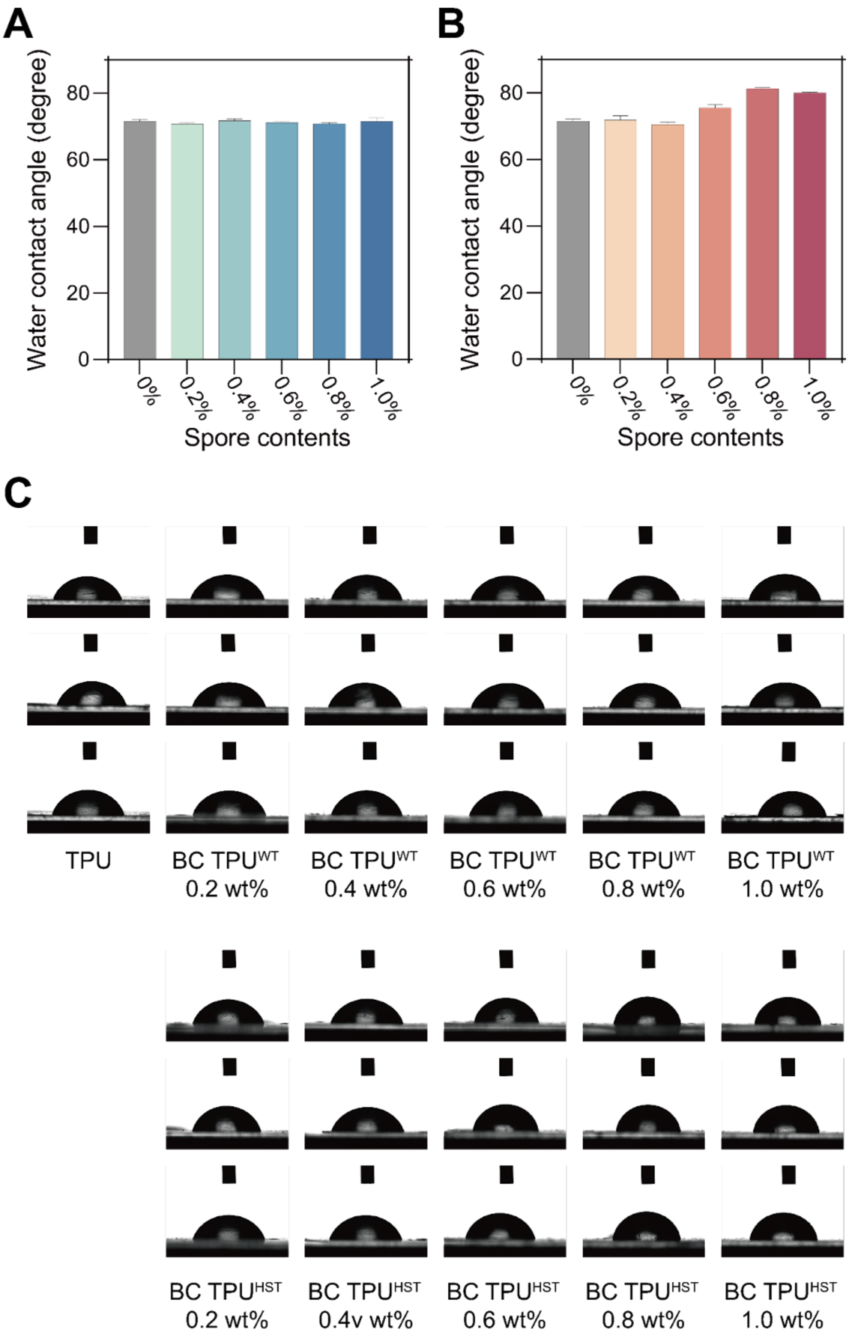

**Supplementary Fig. 8.** Water contact angle of BC TPU$^{WT}$ (A) and BC TPU$^{HST}$ (B). Error bars indicate standard deviations from three independent experiments. (C) Water contact angles were obtained by analyzing the photographs of water droplets on flattened BC TPU$^{WT}$ and BC TPU$^{HST}$.

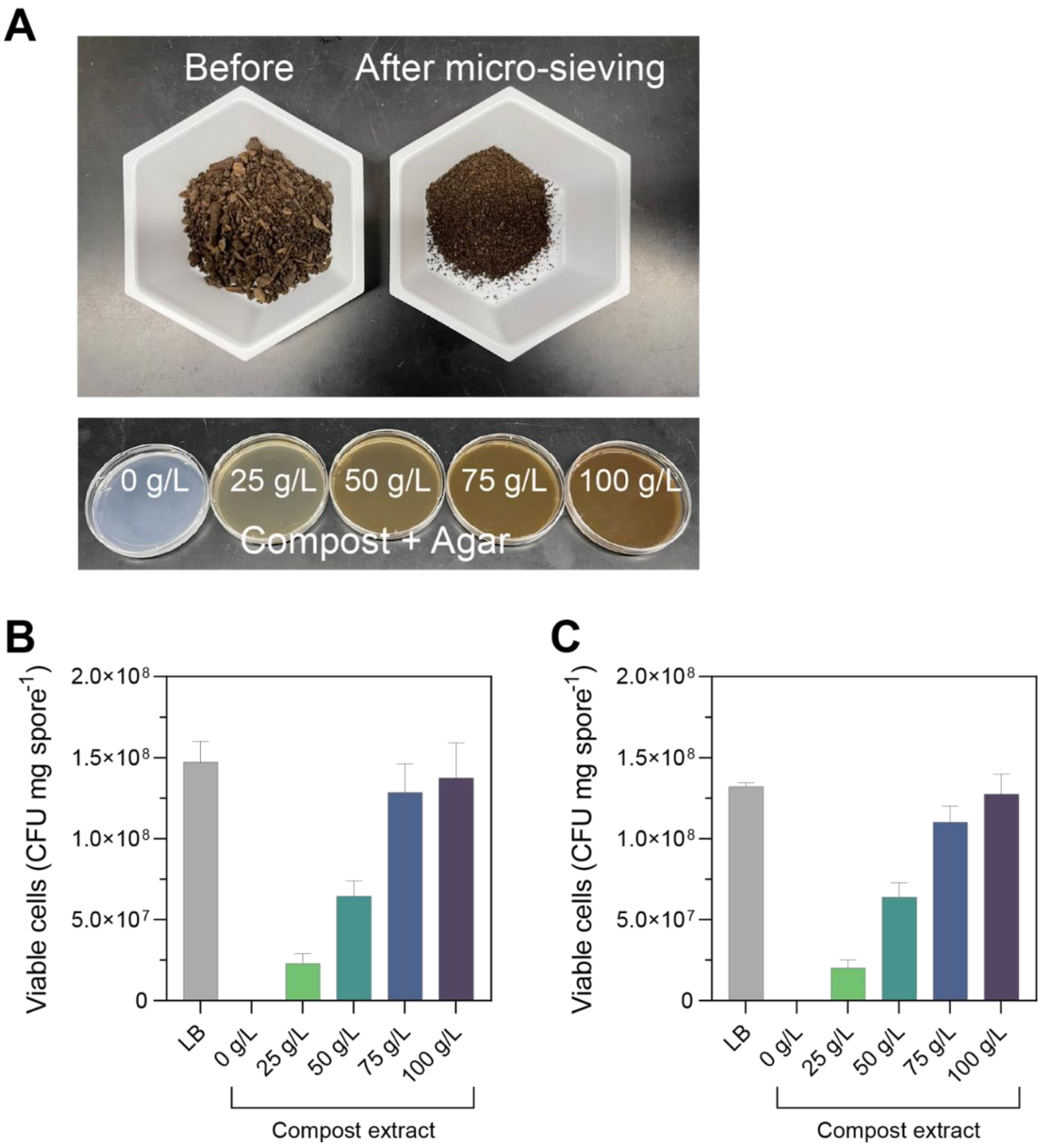

**Supplementary Fig. 9.** (A) Preparation of compost extract gels to assess nutrient availability for spore germination. CFU assays with WT (B) and HST (C) spores on compost gel plates. Error bars indicate standard deviations from three independent experiments.

### Supplementary Note 1: Spore germination on compost extract gels

To confirm *B. subtilis* spores can be germinated by utilizing nutrients in compost, CFU assay for ATCC 6633 WT and HST spores was carried out by using compost gel plates. Compost gel plates were prepared by the following procedure. Dried compost was sieved using a 35 standard mesh screen. The resulting compost powder was suspended in deionized water at various concentrations (25 - 100 g/L). Agarose powder was added to the compost suspension 15 g/L final concentration. The mixture solution was then autoclaved at 121 °C for 20 min. After autoclaving, the settled compost powder was separated, and the supernatant was carefully collected. The supernatant was aliquoted and solidified in petri-dishes.

CFU on LB gel plates served as positive controls. The WT and HST spores showed $1.47 \times 10^8$ and $1.32 \times 10^8$ CFU/mg, respectively, on nutrient-enriched LB plates (**Supplementary Fig. 9**). The CFU of spores on compost gel plates increased with the concentration of compost extract and showed up to $1.37 \times 10^8$ and $1.27 \times 10^8$ CFU $mg^{-1}$, respectively. These values corresponded to 93.3% and 96.4% germination efficiency compared to their positive controls, respectively. This result indicated that compost contains enough nutrients to trigger the spore germination, which is not surprising given the adaptability, fast growth rate, and widespread presence of *Bacillus* sp. in soil[1].

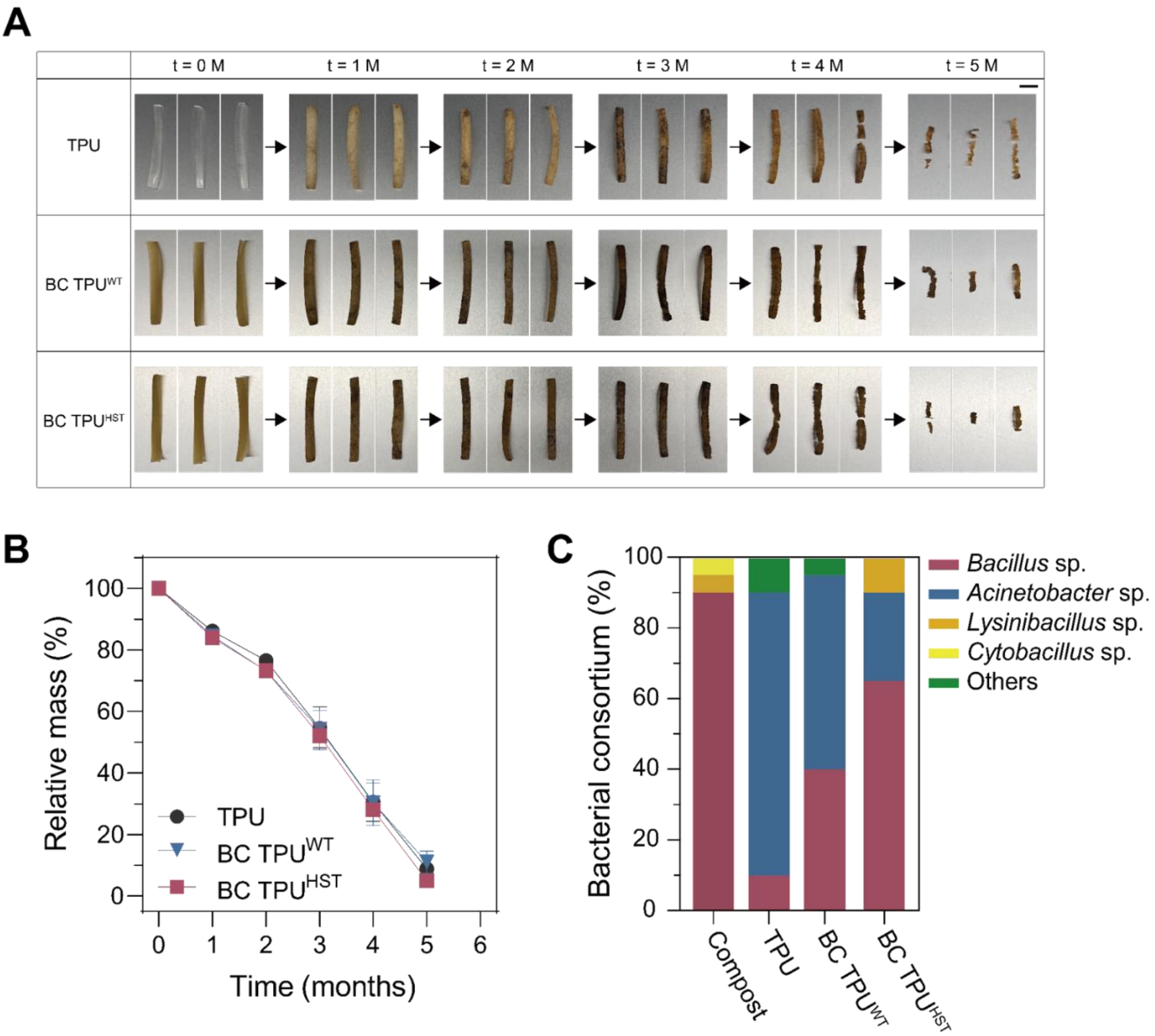

**Supplementary Fig. 10.** (A) Photographs of disintegrated TPU samples during 5 months of incubation in untreated compost at 37°C with a 45-55% of relative humidity (scale bar: 1 cm). Biocomposite TPUs were prepared at 0.8 w/w% spore loading. Mass loss profile (B) and surface microorganism consortium. Error bars indicate standard deviations from three independent experiments. (C) Bacterial consortium of TPU, BC $TPU^{WT}$ and BC $TPU^{HST}$ incubated in untreated compost.

**Supplementary Note 2: Disintegration of biocomposite TPU in untreated compost**

After 5 months of incubation in a rich compost with confirmed microbial activity, TPU, BC $TPU^{WT}$ and BC $TPU^{HST}$ lost 91.1%, 89.0%, and 95.0% of their initial masses, respectively (**Supplementary Fig. 10A-B**). Even though all the TPU samples showed significant mass loss in compost, there was no notable difference in the disintegration rates among TPU and BC TPUs when other microbes were known to be present in the rich compost condition (i.e., a microbially active environment). The respirometry evaluation showed that TPU samples were metabolized to $CO_2$ by the microorganism consortia in the compost (**Supplementary Fig. 11**).

Sequencing analysis revealed that even though the degradation rates of TPU, BC $TPU^{WT}$ and BC $TPU^{HST}$ were similar, the bacterial consortium involved in the degradation of each TPU material was different (**Supplementary Fig. 10C**). *Acinetobacter sp.* in untreated compost was primarily responsible for the biodegradation of pristine TPU, but the portion of *Bacillus sp.* in the microbial consortium was significantly increased on BC TPU samples. It depicts that *B. subtilis* showed comparable disintegration activity toward TPU with the predominant TPU degrader in the compost, *Acinetobacter sp.*. The change of bacterial consortium by the introduction of *B. subtilis* spores into TPU is not surprising because *B. subtilis* is well known environmentally-friendly biocontrol agent, which competes with the pathogenic bacteria in soil and suppress their growth[2]. The improved viability of HST spores, when compared to WT, after melt processing allowed *B. subtilis* to better compete against *Acinetobacter sp.* in compost.

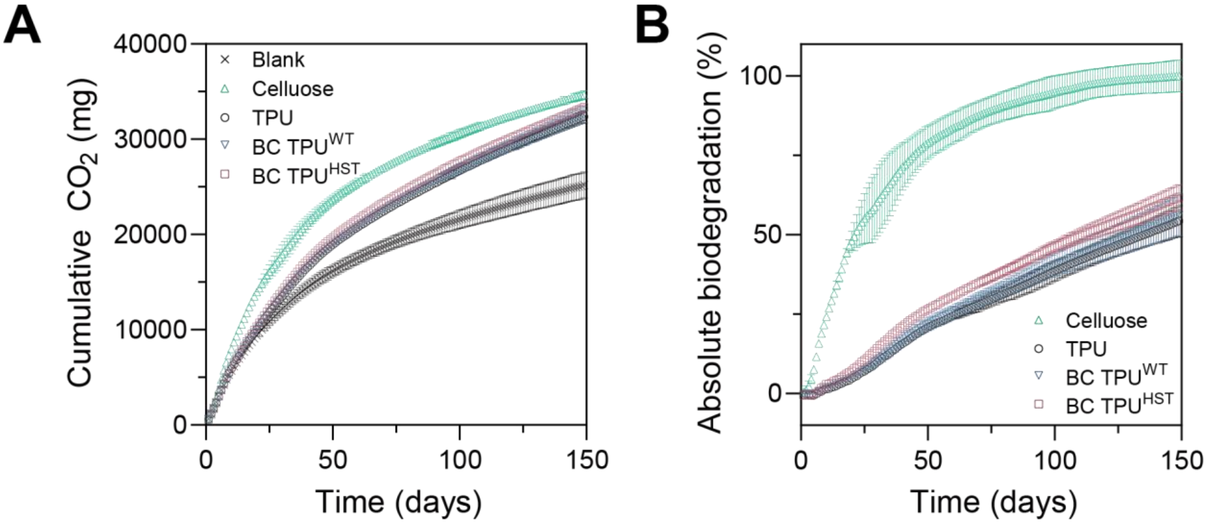

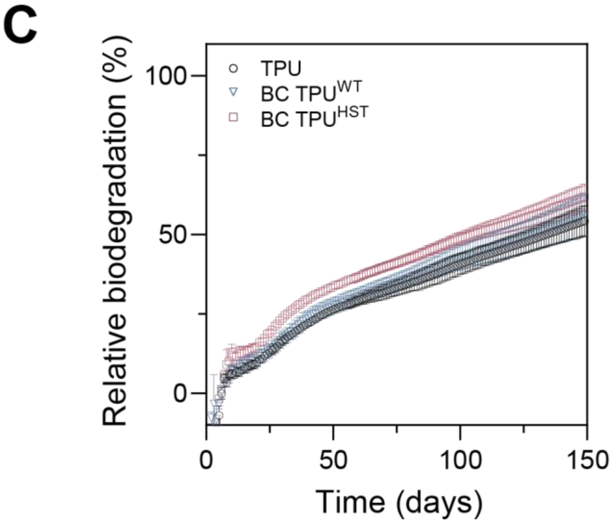

**D**

| | Cumulative $CO_2$ (mg) | Absolute biodegradation (%) | Relative biodegradation (%) |
|---|---|---|---|
| Blank | 25168.5 | N/A | N/A |
| Cellulose | 34640.8 | 100.1 | N/A |
| TPU | 32359.5 | 54.3 | 54.3 |
| BC TPU$^{WT}$ | 32628.5 | 56.3 | 56.2 |
| BC TPU$^{HST}$ | 33334.6 | 62.0 | 61.9 |

**Supplementary Fig. 11.** Cumulative $CO_2$ production (A), absolute biodegradation (B) and relative biodegradation (C) profiles of blank, cellulose, TPU, BC TPU$^{WT}$ and BC TPU$^{HST}$ in compost at 42 °C at 45–60% relative humidity. Error bars indicate the standard deviations from three independent experiments. (D) 149-day averages of cumulative $CO_2$ production and percent biodegradation values for each test group.

**Supplementary Note 3: Respirometry evaluation of TPU**

As a result, TPU, BC $TPU^{WT}$ and BC $TPU^{HST}$ displayed similar mineralization by the end of testing with 25445 mg, 25735 mg and 26441 mg cumulative $CO_2$ production, respectively, after 149 days of incubation. Absolute biodegradation of TPU, BC $TPU^{WT}$ and BC $TPU^{HST}$ was calculated as 54.3%, 56.3% and 62.0%, respectively. Given that the absolute biodegradation of cellulose was 100.1%, relative biodegradation of TPU, BC $TPU^{WT}$ and BC $TPU^{HST}$ corresponded to 54.3%, 56.2% and 61.9%, respectively (**Supplementary Fig. 11**).

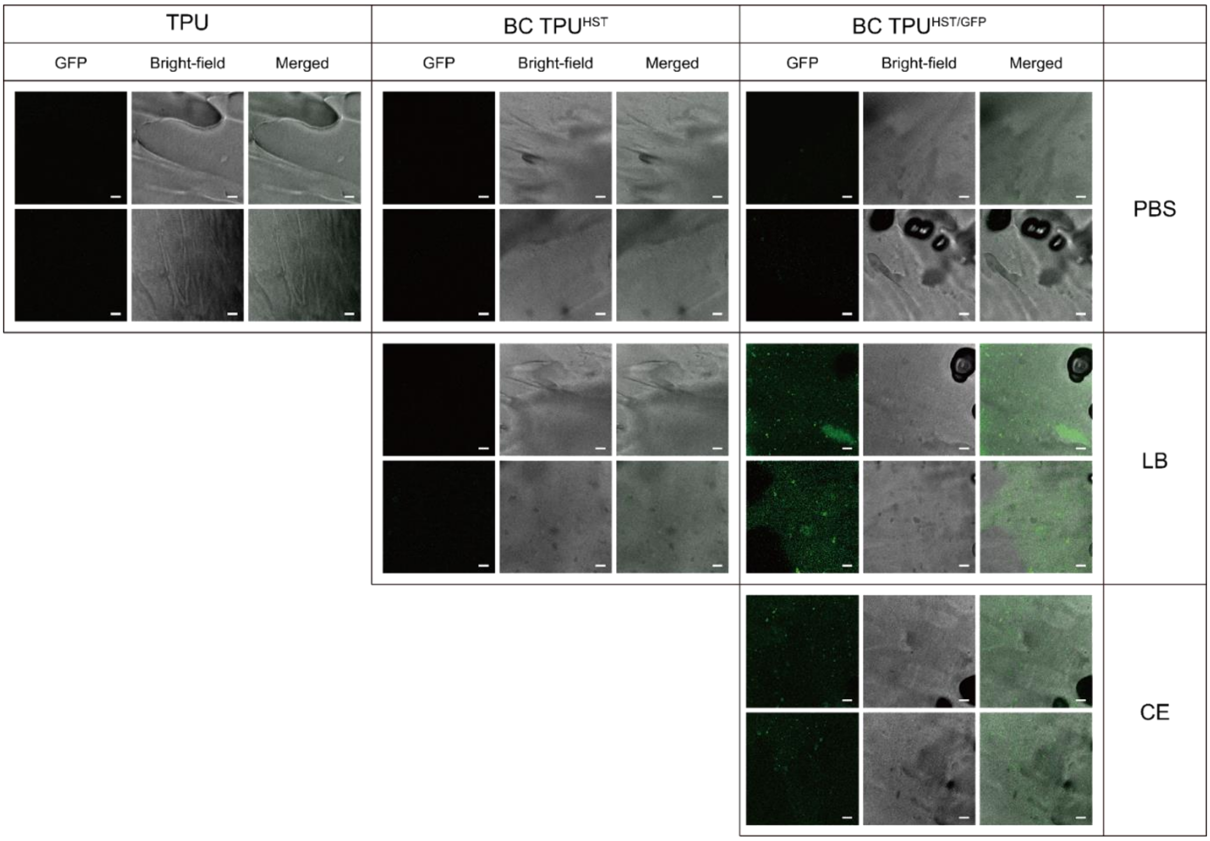

**Supplementary Fig. 12.** Fluorescence, bright-field and merged images (left to right) of TPU, BC TPU$^{WT}$ and BC TPU$^{HST}$ incubated in PBS, LB or CE obtained by CLSM. Scale bars: 10 µm.

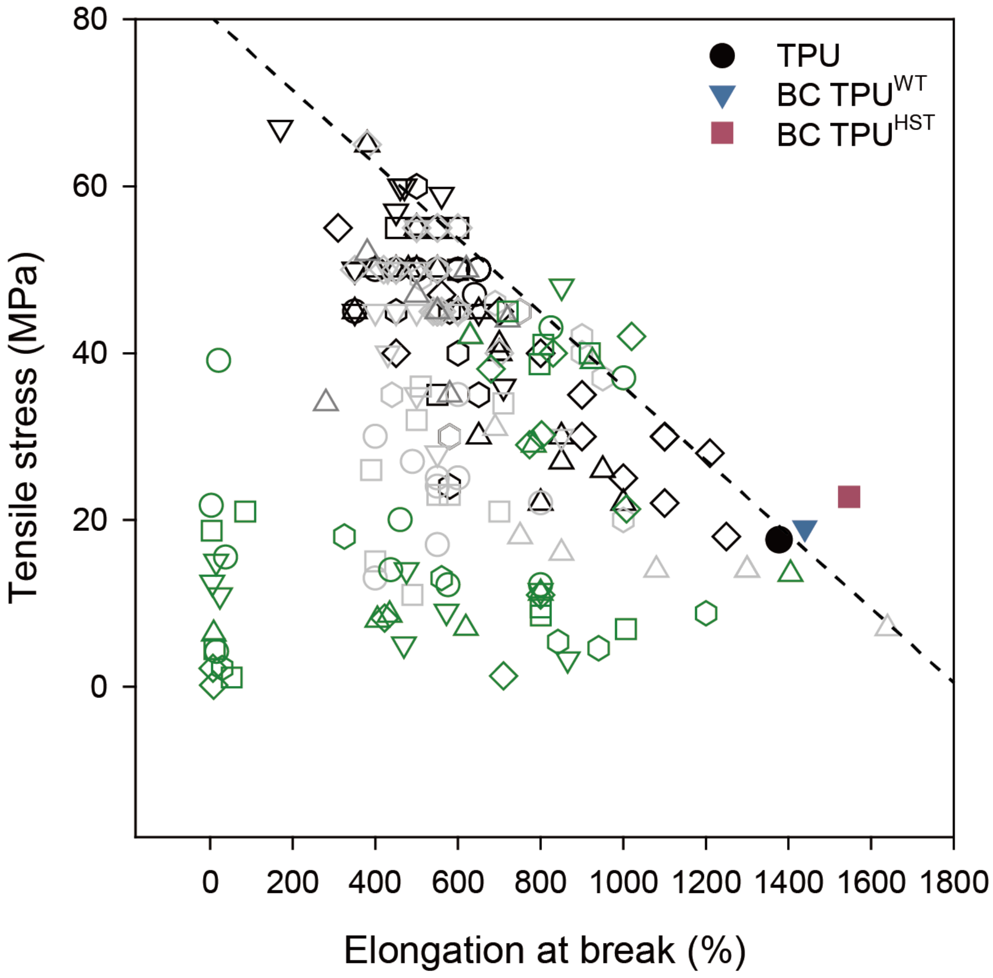

**Supplementary Fig. 13.** Elongation at break and tensile stress of commercially-available TPUs (grayscale) were plotted with BC TPUs (colored) developed in this work. Previously reported biodegradable TPUs are plotted with green symbols[3–8]. Tensile properties of commercially-available TPUs were adapted from the BASF Elastollan® product range. The trade-off barrier (dashed line) was plotted by using the tensile properties of top 15 commercial TPUs that mutually exhibit high tensile stress and elongation at break.

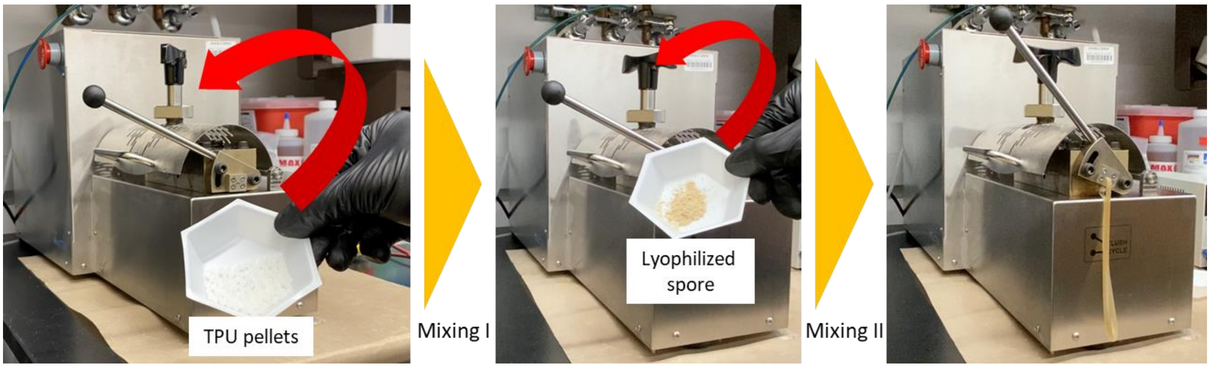

**Supplementary Fig. 14.** Fabrication of BC $TPU^{HST}$ using benchtop twin screw extruder.

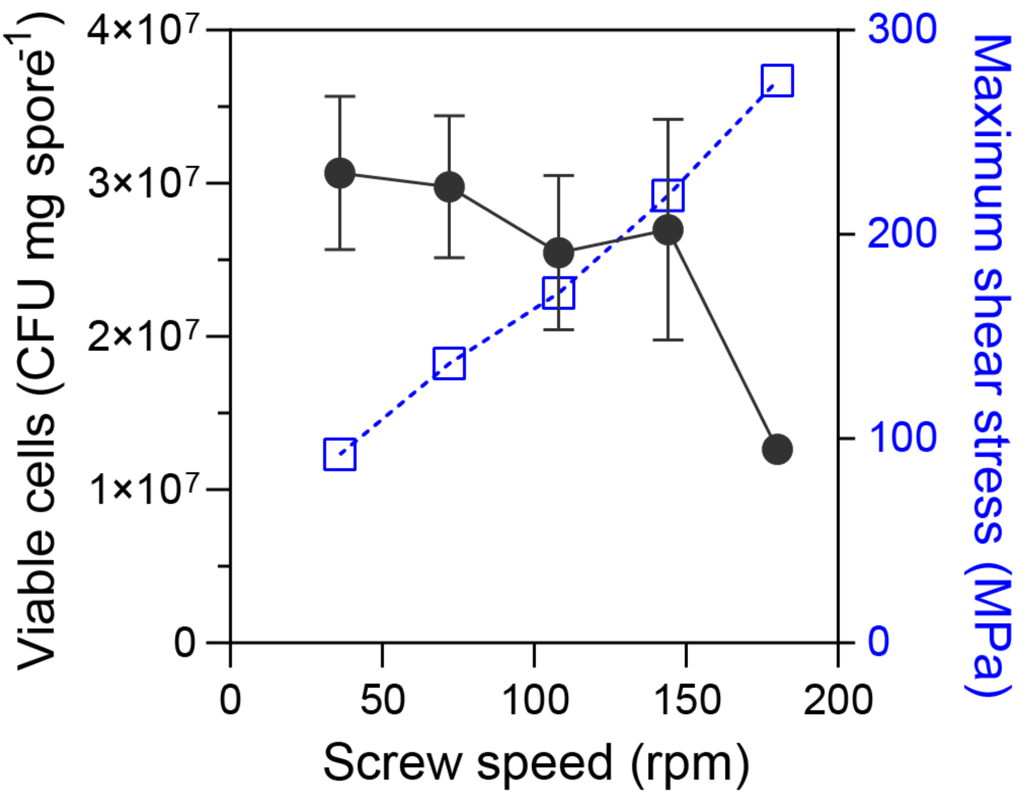

**Supplementary Fig. 15.** Spore viability and maximum shear stress at different screw speeds.

**Supplementary Table 1.** Number of viable cells in untreated and autoclaved compost. Cell viability of autoclaved compost was also tested after 5 M of incubation under 37 °C at 45-55 % relative humidity.

| **Sample** | **Viable cells (CFU / mg compost)** | **Cell survivability (%)** |
|---|---|---|
| Compost | 1.6 (±0.3) x $10^6$ | 100 |
| Autoclaved compost | 2.6 (±0.3) | 0.0002 |
| Autoclaved compost after 5 M of incubation | 7.9 (±0.1) x $10^3$ | 0.5 |

**Supplementary Table 2.** Elemental metal analysis of compost performed via total acid digestion.

| **Element** | **Recommended specification** | **Compost for gravimetric biodegradation** | **Compost for respirometry** |
|---|---|---|---|
| Al (aluminum) | No limit | 14700 | 11545 |
| As (arsenic) | < 20 ppm | 3.25 | 2.31 |
| B (boron) | No limit | 25.08 | 31.30 |
| Ca (calcium) | No limit | 15233 | 26901 |
| Cd (cadmium) | < 2 ppm | 0.18 | 0.88 |
| Cr (chromium) | < 100 ppm | 19.19 | 17.60 |
| Cu (copper) | < 100 ppm | 101.26 | 64.50 |
| Fe (iron) | No limit | 11677 | 11421 |
| K (potassium) | No limit | 6033 | 8758 |
| Mg (magnesium) | No limit | 2695 | 3910 |
| Mn (manganese) | < 3500 ppm | 532.9 | 613.0 |
| Mo (molybdenum) | < 440 ppm | 2.01 | 2.31 |
| Na (sodium) | No limit | 1154 | 1224 |
| Ni (nickel) | < 50 ppm | 7.76 | 4.79 |
| P (phosphorus) | No limit | 11578 | 10471 |
| Pb (lead) | < 75 ppm | 14.84 | 23.80 |
| S (sulfur) | No limit | 3413 | 2920 |
| Total Solids (%) | 50 – 55% | 59.05 | 30.5 |
| Volatile Solids (%) | N/A | 40.03 | 26.78 |

| | | | |
|---|---|---|---|
| Ash (%) | <70% | 35.42 | 3.72 |
| pH | 7.0 – 8.2 | 7.02 | 7.40 |
| Carbon (%) | N/A | 20.28 | 27.04 |
| Nitrogen (%) | N/A | 1.74 | 2.49 |
| C:N | 10 – 40 | 11.66 | 10.86 |

**Supplementary Table S3.** Carbon and nitrogen content of test materials and positive control for respirometry composting experiment. Limit of detection is 0.10%.

| **Sample** | **Total Carbon (%)** | **Total Nitrogen (%)** |
|---|---|---|
| Cellulose | 42.88 | 0.19 |
| TPU | 58.85 | 7.36 |
| BC $TPU^{WT}$ | 58.76 | 7.28 |
| BC $TPU^{HST}$ | 59.37 | 7.26 |
